# Patterns of Arc mRNA expression in the rat brain following dual recall of fear- and reward-based socially acquired information

**DOI:** 10.1101/2021.12.30.474464

**Authors:** Laura A. Agee, Emily N. Hilz, Dohyun Jun, Victoria Nemchek, Hongjoo J. Lee, Marie-H. Monfils

## Abstract

The ability to learn new information and behaviors is a vital component of survival in most animal species. This learning can occur via direct experience or through observation of another individual (i.e., social learning). While research focused on understanding the neural mechanisms of direct learning is prevalent, less work has aimed at understanding the brain circuitry mediating the acquisition and recall of socially acquired information. We aimed to further elucidate the mechanisms underlying recall of socially acquired information by having rats sequentially recall a socially transmitted food preference (STFP) and a fear association via fear conditioning by-proxy (FCbP). Brain tissue was processed for mRNA expression of the immediate early gene (IEG) *Arc*, which reliably expresses in the cell nucleus following transcription before migrating to the cytoplasm over the next 25 minutes. Given this timeframe, we were able to identify whether *Arc* transcription was triggered by STFP recall, FCbP recall, or following recall of both memories. Surprisingly – and contrary to past research examining expression of other IEGs following STFP or FCbP recall separately – we found no differences in any of the *Arc* expression measures across a number of prefrontal regions and the vCA3 of the hippocampus between controls, demonstrators, and observers, though we did detect an overall effect of sex in a number of regions. We theorize that these results may indicate that relatively little *Arc*-dependent neural restructuring is taking place in the prefrontal cortices following recall of a recently socially acquired information or directly acquired fear associations in these areas.

## Introduction

An animal’s capacity to survive in a new environment is largely contingent on their ability to learn about and adapt to their surroundings by identifying both potential threats and sources for fulfilling essential needs. Humans, perhaps more than any other species, are particularly adept at acquiring new strategies to deal with environmental challenges or exploit avenues for securing resources. One of the primary ways in which we are able to learn such strategies at an individual level is through receiving instructions or observing an experienced individual, i.e., via social learning. As such, it should not be surprising that deficits in the ability to socially learn have the potential to significantly impair functioning. This can be seen in autism spectrum disorders, in which much of the symptomology is thought to arise from impairments in the social attention/reward systems and, by extension, the social learning system [1–3].

Conversely, there are also drawbacks if social learning occurs too indiscriminately. While valuable information and adaptive behaviors can be acquired socially, this does not preclude individuals from socially acquiring false information or maladaptive behaviors through the same pathway. Clinically, this is often seen in phobias, which are commonly reported to have been acquired through observation or instruction (e.g., watching a parent react with extreme fear to a spider or receiving dire warnings about the danger of spiders, respectively) rather than by direct experience [4,5]. Socially acquired phobias may also be disruptive in ways directly acquired phobias are not, because the individual has not directly experienced the aversive consequences in relation to the feared stimuli. As such, they are free to imagine an associated outcome that may be more intense than what occurs in reality. In line with this idea, individuals with socially acquired phobias report increased cognitive symptomology [6] and respond more favorably to certain treatment methods [4] than do individuals with directly acquired phobias.

To truly understand and subsequently develop optimal treatments for conditions arising from under- or over-performing social learning, a thorough understanding of the brain mechanisms that underlie the social learning process is an essential first step. One of the primary methods we have for exploring such mechanisms are non-human animal models. In rodent species, fear-based social learning has been demonstrated to occur under multiple conditions, including: (1) context or stimulus associated fear acquired by observation through a barrier of a conspecific experiencing pain in a novel environment or following the presentation of a novel stimulus [7,8], (2) enhanced acquisition of natural behaviors by observation of a conspecific responding to a threatening stimuli [9–11], and (3) by observation of a fear conditioned demonstrator reacting to the fear-associated stimuli post-conditioning in a paradigm known as fear conditioning by-proxy (FCbP) [12–14].

While similar reward-based models of social learning in rodents have proven somewhat more difficult to develop [15], one reliable and well-established model of reward-based socially mediated learning does exist in the social transmission of food preference (STFP) paradigm [16–19]. In the STFP paradigm, rats assigned to the ‘demonstrator’ condition consume a novel food (generally powdered chow mixed with flavoring, such as cinnamon) before interacting with a naïve rat assigned to the ‘observer’ condition. When observers are later given the choice to consume either the demonstrated flavor or an entirely novel flavor, they reliably show the tendency to consume more of the demonstrated flavor. This effect has been shown to be mediated by the semiochemical carbon disulfide (CS2) which is present in the nasal cavity of rats and, when paired with a novel scent, is sufficient to induce a preference for similarly scented foods [17].

In rodents, there has been a fair amount of research examining the brain mechanisms mediating the acquisition and recall processes for the social transmission of food preference task [20–24] and, to a lesser extent, socially acquired fears [7,12,13,25–27]. Results from research into the latter topic have also found that there are a number of brain areas that seem to be uniquely activated during social fear learning and not direct fear learning [13,27]. Furthermore, integrative models considering the results from both human and non-human animal research into the brain circuitry underlying the social learning of appetitively and aversively motivated behaviors/associations posit that, while there does seem to be considerable overlap between the brain areas governing direct learning processes and social learning processes, activity in some unique brain regions is required for social learning to occur [13,28].

While the neural mechanisms involved in the social acquisition of tasks and information has received some exploration, research explicitly comparing the storage of memories acquired by social learning to memories acquired by direct learning is, to our knowledge, almost nonexistent. In the experiment described here, we attempted to examine activation in various brain regions following recall of a socially acquired memory from both a reward- and fear-based task. Rats were trained in a reward-based form of social learning, STFP, and a fear-based model of social learning, FCbP, after which we initiated sequential recall of both memories. The tissue from these rats was then processed for mRNA expression of the immediate-early gene (IEG) – a class of genes which are rapidly transcribed following neuronal firing or other cellular stimuli - *Arc* which, when transcribed, produces the mRNA for the activity-regulated cytoskeleton associated (Arc) protein. Arc mRNA has a predictable pattern of expression such that in the first 5 minutes following transcription it is expressed in the nucleus of the cell and, after about 25 minutes, migrates to the cytoplasm surrounding the nucleus [29]. As such, cells stained for *Arc* mRNA that are activated at both timepoints show expression in both the cytoplasm and nucleus, allowing for precise localization of cell populations activated in multiple tasks. By analyzing the expression of *Arc* mRNA in rat brains perfused following the sequential recall of FCbP and STFP tasks, we aimed to identify brain regions uniquely involved in retrieval of socially acquired information. The anterior cingulate cortex [7,13,28], orbitofrontal cortices [23,24], and infralimbic and prelimbic cortices [23,30–32] were all of particular interest given past research which has implicated them in fear learning, social fear learning, STFP learning, or some combination of the three.

## Methods

### Subjects

Subjects were male and female Sprague-Dawley rats bred in house in the Animal Resource Center of the University of Texas at Austin. Eight breeding pairs were used to produce the subjects for Cohort 1 of this experiment - with seven successfully breeding - while eight separate breeding pairs were used to produce the subjects for Cohort 2. Female breeding animals were Sprague-Dawley rats (between 215-260g at arrival) obtained from Charles-Rivers (Wilmington, MA, USA) while male breeding animals were Sprague-Dawley rats (most between 275-300g at arrival with one at 230g) obtained from Harlan (now Envigo) (Houston, TX, USA) to prevent accidental inbreeding. All rats were paired off with an opposite-sex cage mate following arrival to the colony. Once a female began to show clear signs of pregnancy, her paired male was removed from the cage and rehoused.

Once delivered, pups were weaned into triads of same-sex siblings at post-natal day 21 (P21) to help ensure social fear learning [26]. Spare pups were weaned into triads or dyads with unrelated rats and used in other experiments at the University of Texas at Austin. Female pups from our second cohort litter were used in other experiments. The final number of pups used for this experiment were n = 27 for Cohort 1 females, n = 36 Cohort 1 males, and n = 27 Cohort 2 males. Pups being utilized in this experiment were allowed to mature with minimal disturbances aside from routine animal husbandry procedures (e.g., cage changes) until habituation procedures (Females triads) or dominance assessment procedures (Male triads) began. All Cohort 1 rats started on habituation procedures between P106-P112 days of age (young adulthood) and all Cohort 2 rats were started between P99-P118 days of age. All subjects were kept on a 3 pm – 3 am lights off light-cycle and all experimental procedures were completed during the subjects’ dark cycle. All parts of this experiment were conducted in compliance with the National Institutes of Health Guide for the Care and Use of Experimental Animals and were approved for use by The University of Texas at Austin Animal Care and Use Committee.

### Apparatus and Stimuli

#### Fear Conditioning

All fear conditioning and fear conditioning by-proxy procedures were completed in standard conditioning chambers (30.48 cm x 25.4 cm x 30.48 cm) constructed of clear plexiglass walls in the front and back, two steel walls on the side, and a plexiglass ceiling with a hole in the center. The flooring of the chamber was a row of stainless-steel rods connected to a shock generator (Coulbourn Instruments, Allentown, PA). All chambers were enclosed in acoustic isolation boxes (Coulbourn Instruments) and lit with an internal red light. Behavior was recorded by closed-circuit cameras (Panasonic™ WV-BP334) mounted above the conditioning chambers with the lens inserted through the hole in the plexiglass ceiling. Chambers were fully wiped down with 70% alcohol solution between each subject. All stimulus delivery was controlled using the Freeze Frame software (Coulbourn Instruments). The conditioned stimulus (CS) was a 20 second tone (5kHz, 80 dB) and, in procedures with multiple CS presentations, a variable inter-trial interval (ITI) averaging 180 seconds. The unconditioned stimulus (US) was a 1 mA shock that was 500 milliseconds in duration and co-terminated with the conditioned stimulus.

#### Social transmission of food preference

All STFP procedures took place in a room adjacent to the room containing the conditioning chambers. Novel diets were composed by mixing 100g of powdered 5LL2 Purina rodent chow with either 1g of McCormick ground cinnamon (diet Cin) or 2g of Hershey cocoa powder (diet Co). The Plain diet, which was given to all rats during the food restriction period and to Control rats on the terminal day of experimental procedures, was unadulterated powdered 5LL2 Purina rodent chow. All powdered chows – both during food restriction and experimental procedures – were presented in hanging food cups that were constructed from 4 oz. glass jars and 12-gauge steel utility wire. Food cups were rinsed then wiped down with a 70% ethanol solution before being washed thoroughly with soap and water between every use. All consummatory phases of the STFP experimental procedures took place in standard rat cages (26.7 cm x 48.3 cm x 20.3 cm), with every animal receiving a fresh cage. The interaction phase (STFP acquisition phase) took place in a large plastic bin (50.5 cm × 39.4 cm × 37.5 cm) with wood chip floor bedding that was replaced between every group. Plastic bins were wiped down thoroughly with Windex between each session.

### Overview of Experimental Design & Social Learning Procedures

(See Fig 1 for a graphical overview)

**Fig 1.**
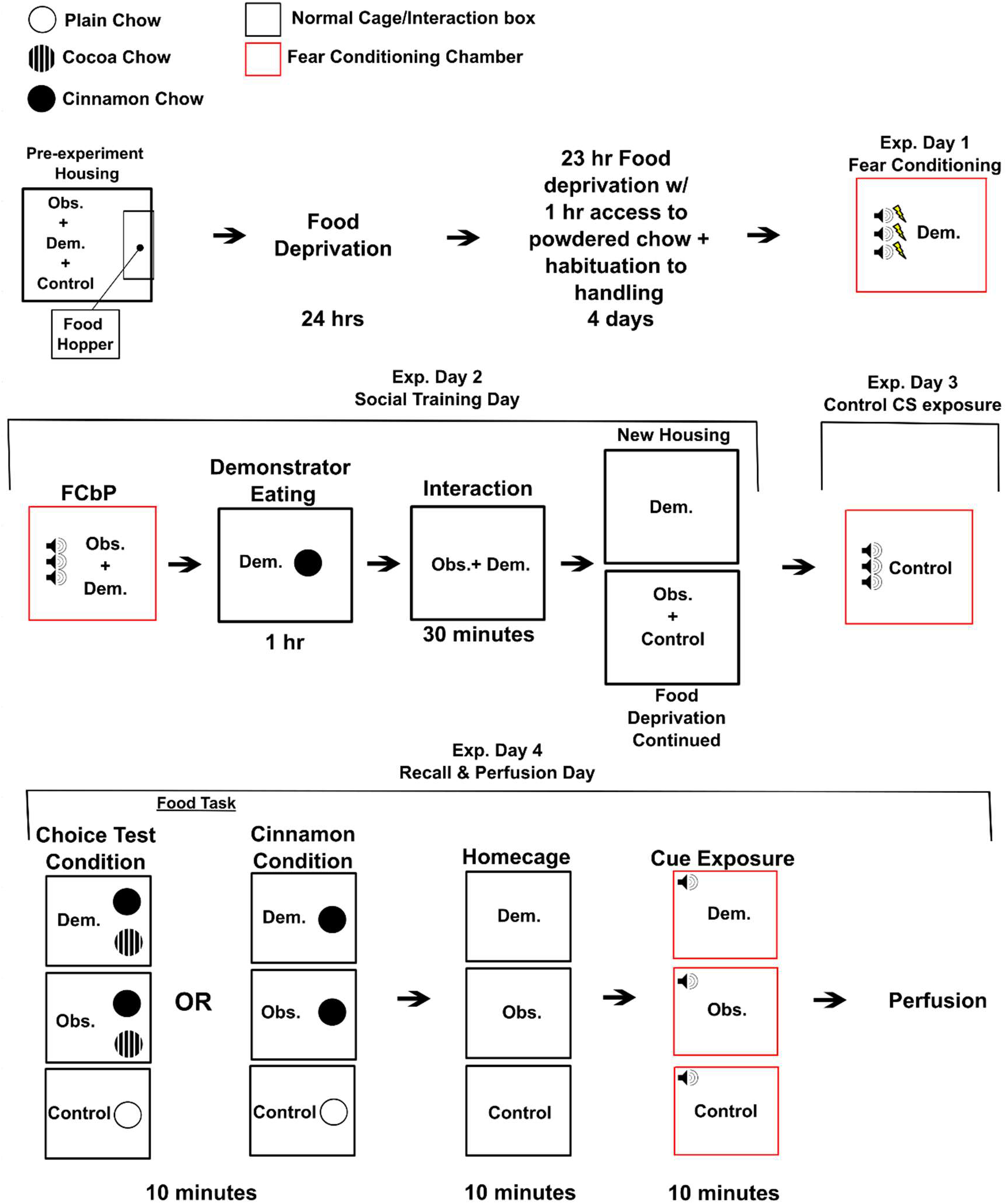
Overview of Experiment Design. This figure outlines the treatment of rats on each day of the experiment from the first day of food restriction on.

All rats were food restricted for five days and habituated to handling and the room where STFP procedures would take place for four days immediately prior to day 1 of experimental procedures. While habituation procedures ended prior to day 1 of experimental procedures, food restriction continued through to the end of the experiment. One animal from each triad of rats was assigned to one of three conditions: Demonstrator, Observer, or Control. Cohort 2 male triads had been assessed for dominance and all showed a clear hierarchy and were assigned such that the dominant rat was the Demonstrator and a subordinate was the Observer to enhance social transmission of fear [13]. Individual triads were further randomly subdivided into groups where the Demonstrator and Observer would receive a choice test at STFP recall (Choice) and groups where they would receive only the demonstrated food (Cin).

On day 1 of the experimental procedure, rats assigned to the Demonstrator condition were moved to conditioning chambers and allowed to habituate for 10 minutes before they were exposed to 3 CSs that co-terminated with a painful shock (see **Apparatus and Stimuli** for specifics). Following fear conditioning procedures, Demonstrators were moved back to their original home cage. On day 2 of experimental procedures, 24 hours after fear conditioning, Demonstrators were returned to the conditioning chambers with their cage-mate assigned to the Observer condition and put through the FCbP procedure. Immediately following the FCbP procedure, Observer rats were returned to their home-cage while Demonstrators were moved to an adjacent room and given 1 hour to consume powdered chow flavored with cinnamon. After an hour had passed, Observers were moved to an interaction bin with their Demonstrator and allowed to interact with them for 30 minutes to allow for acquisition of a socially transmitted food preference. Previous research from our lab has validated these timepoints as being sufficient for STFP transmission [33]. Afterwards, Observer rats were returned to their home-cage while Demonstrator rats were moved to single housing to prevent further STFP transmission to the Observer or Control. On day 3 of experimental procedures, Control rats were moved to conditioning chambers alone and, following 10 minutes of habituation to the chamber, were presented with three 20 second CSs with no accompanying shock. This was done on a separate day to minimize the possibility of lingering alarm pheromones – which are known to be released by rats in response to threatening stimuli and effect conspecific learning [34] – still being present in the chamber.

On the terminal day of the experiment, day 4, recall was initiated for both the socially transmitted food preference and the fear conditioning/fear conditioning by-proxy memories. All Observers and Demonstrators from triads assigned to the Choice condition were allowed 10 minutes *ad libitum* access to both cinnamon and cocoa flavored diets, while Observers and Demonstrators from triads assigned to the Cin condition were given 10 minutes *ad libitum* access to the cinnamon diet only. In all triads, Control rats were given 10 minutes *ad libitum* access to plain powdered chow. Immediately after this, rats were returned to their home-cage and left undisturbed for a 10-minute period before being moved back to the lab space and being placed in the conditioning chambers. All rats were then given a 3-minute habituation period to the chamber before being presented with a single 20 second CS. 5 minutes after the end of the CS, all rats were euthanized via injection of a pentobarbital and phenytoin solution (Euthasol; Virbac Animal Health) and perfused. Their brains were later processed for *Arc* mRNA expression. Given the time course of our terminal procedure and the known migration timeframe of *Arc* mRNA [28], increases cytoplasmic expression of *Arc* mRNA would be due to STFP recall procedures, while nuclear expression would be due to FC/FCbP recall procedures, with cells showing dual activation having been activated at both timeframes (see Fig 2; also, see **Tissue Analysis** for details on tissue treatment and processing).

**Fig 2.**
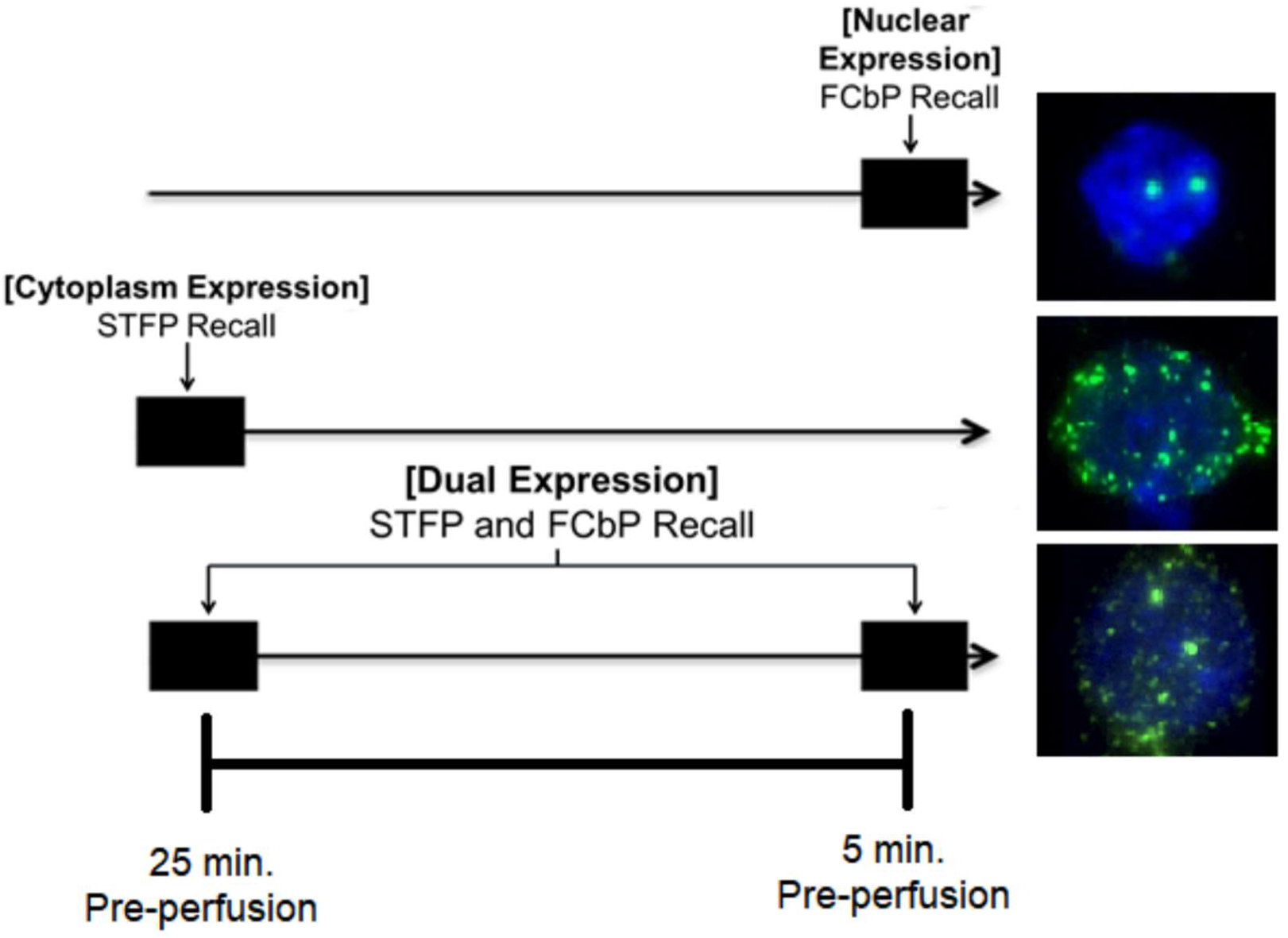
Patterns of *Arc* mRNA Expression. The above figure shows the pattern and area within a cell in which we would see *Arc* mRNA expression triggered by activity at the FCbP recall timepoint, the STFP recall timepoint, or activity that was triggered at both timepoints.

### Procedures

#### Habituation and food restriction

All habituation took place just prior to the first day of experimental procedures. Habituation consisted of each cage of rats being moved into the room in which all STFP experimental procedures would take place and being allowed to habituate to the room for 15 minutes. During this period, each rat was picked up and handled by the experimenter that would be running behavior for 2 minutes to habituate them to handling and that individual. All habituation procedure took place in a dark room under red light, and all rats received 4 days of habituation. Food restriction began the day before habituation began and persisted to the end of the experiment. At the start of food restriction, the food pellets that all subjects had been eating were removed from the cage. Subsequently, all cages were given daily *ad libitum* access to a hanging jar full of plain, powdered Purina 5LL2 diet in their home-cage. Rats were weighed daily starting at the beginning of food restriction until the experiment was over to ensure no unusual loss in weight.

#### Play behavior dominance assessment

A day prior to play dominance assessments, all males were moved to single housing to promote social play behavior. Following a 24-hour isolation period, individuals from each triad were moved to a large plastic bin (50.5 cm × 39.4 cm × 37.5 cm) with woodchip bedding and a camera mounted overhead to record behavior. Rats were allowed to interact for 15 minutes before being removed from the box and returned to single housing. This was repeated for 3 sessions, after which rats were returned to their triads and left undisturbed until the start of the milk competition dominance assessment. Behavior was scored as described below, and rats in Cohort 2 were assigned to one of three dominance ranks based on their behavior as following with past research on dominance hierarchies in rats [35]: Dominant, Subordinate 1, or Subordinate 2. Male rats in cohort 1 were randomly assigned condition regardless of dominance rank. As described in Jones & Monfils [13], dominant rats were the rats that received most nape contact (i.e., play initiations), while subordinate 1 was the rat that initiated the dominant the most, and subordinate 2 tended to be avoidant. While all Cohort 1 males were used, male triads in Cohort 2 that did not show dominance hierarchies were removed from the study and used in other experiments. Analysis of play behavior dominance included only Cohort 2 rats.

#### Milk competition dominance assessment

In order to validate dominance assignments made using the play behavior assessment, we recorded and scored the behavior of male rats allowed access to a desired resource (sweetened milk solution), a dominance assessment that our lab previously found to be effective [13]. The milk solution used in this dominance assessment was a mixture of 2/3 tap water and 1/3 sweetened condensed milk (Eagle™) stored in a 2 oz glass jar filled to the top with the solution. Prior to running the dominance assessment, rats from all male triads were moved to single-housing and given access to a full jar of the milk for 5-hours to ensure that each individual rat had the opportunity to overcome their neophobia of the milk solution. Following this, rats were returned to their triads and allowed access to a full jar of the milk solution as a group daily for four days. In order to assure that rats would be motivated to drink, food hoppers were removed from all triads 12 hours before the milk was introduced. Following the 3-hour milk access period, hoppers were returned until removal time for the next day of habituation.

Once habituation to the milk solution had been completed, triads were run through the formal dominance assessment. As during habituation, food hoppers were removed 12 hours before the start of assessment to promote competition. 2 oz glasses were filled with to the top with the milk solution and secured with adhesive strips to the bottom of a large plastic bin (50.5 cm × 39.4 cm × 37.5 cm) with woodchip bedding and portable cameras were mounted above the box for an over-the-head view of all behavior. Rats were placed in the bin and allowed access for either 12 minutes (Cohort 1) or 10 minutes (Cohort 2) before being removed and returned to their triads. While only two sessions of the competition were run for our Cohort 1 males, three sessions were run for Cohort 2 in an attempt to obtain clearer dominance hierarchies.

### Behavioral Scoring

All behavioral scoring for this experiment was completed using the Behavioral Observation Research Interactive Software (BORIS) [36].

#### Play dominance scoring

Behavior was scored for the full play session, with both offensive play behaviors (i.e., play initiations or attacks) and defensive play behaviors (i.e., response to play initiations, specifically nape contact) being scored (see [13,35,37]). The following offensive play behaviors were scored: (1) Nape contact, contact of a rat’s snout with the nape of another rat and (2) Boxing, which occurred when rats reared and punched at each other with their front legs. The defensive behaviors scored for were: (1) Counter, in which the attacked rat turns to face the attacking animal to launch an attack of their own; (2) Evasion, in which the attacked rat flees from the attacker; (3) Full rotation, in which the target rotates fully into a supine position; (4) Half rotation, in which the targeted animal responds to the attack by shifting their body laterally to break contact without fully losing their feet; (5) No response, in which the target either freezes or carries on at a normal pace in response to attack. The identity of both the initiating rat and their target was noted for every instance of play behavior. Across all sessions, the total nape contacts received for each individual rat was tallied and divided by the total number of nape contact initiated in the cage to determine the percent of contacts each rat had received. If a rat had received a disproportionate amount of contact (>40%) they were deemed the dominant animal.

#### Milk competition scoring

Behavior for milk competition began to be scored as soon as all rats were in the bin and the experimenter had exited the footage. Behavior was scored in 1-minute bins for 10-12 minutes. The duration of each subject drinking from or monopolizing the milk jar (i.e., drinking from or having paws/body on the jar and preventing the other rats’ access) was scored for each 1-minute interval. To calculate percent monopolization of the resource, the total time all rats spent drinking in each bin was summed and the time spent drinking for individual rats was divided by that value. The amount of time spent drinking was then plotted based on play behavior dominance assignments for all rats.

#### Fear conditioning by-proxy social contact scoring

Past research from our lab has indicated that there is a strong relationship between the amount of fear displayed by observers at the long-term memory test and the time spent interacting with their Demonstrator during the CS in males [13] and after the CS in females [14,26]. As such, videos of the social acquisition phase of fear-conditioning by proxy were scored for social interaction between the Observer and Demonstrator for each 20 second period during the CS presentation and the 20 second period immediately following each CS presentation to provide a secondary index of fear acquisition. Social contact was scored when Observer and Demonstrator animals made contact other than in passing during the cue period (during CS contact) or in the 20 seconds following the CS (post CS contact). The percentage of each score period spent in contact with the Demonstrator was calculated. Data for percent contact during the cue period for males and data for the percent contact immediately following the cue period for females was pulled and combined into a single “relevant contact” measure to be used in all final statistical analyses.

#### Choice test scoring

Videos of the choice test to initiate recall of a socially transmitted food preference were scored for the amount of time that a given rat spent interacting with a food cup based on whether it contained the demonstrated/already consumed diet (diet Cin) or the novel diet (diet Co). This was done as a potential secondary measure of food preference, as we anticipated that due to the choice test being abnormally short (10 minutes) by necessity that we might be unable to detect preferences based on amount eaten alone. Interaction with the food cup was scored for whenever a rat was physically in contact with and not actively moving away from the cup (i.e., front paws in contact with the jar, head inside jar, climbing on top of the jar, or actively eating from the jar). For statistical analysis, we calculated the percent of time spent interacting with a cup containing a given diet based on the total amount of time spent interacting with either cup (e.g., for diet Cin, Percent time = Time_Diet Cin_/(Time_Diet Co_ + Time_Diet Cin_)). The full 10-minute choice test session was scored for all rats that underwent the choice test with the exception of one rat whose video was unavailable due to recording equipment failure.

### Tissue analysis

To minimize degradation of mRNA by ribonuclease (RNase), all equipment and surfaces used during brain preparation and processing were sanitized regularly with either RNase AWAY™ (Thermo Scientific; Waltham, MA, USA) or RNAseZap™ (Ambion; Grand Island, NY, USA).

#### Brain Preparation

Immediately following euthanasia, subjects were perfused intracardially using a 4% paraformaldehyde (PFA) solution. Brains were then removed and submerged in the 4% PFA solution to allow post-fixation for 24-48 hours. Once post-fixation was complete, brains were transferred to a solution of 30% sucrose in phosphate buffered saline for cryoprotection. Once brains had sunk to the bottom of the vial, indicating sufficient sucrose uptake for cryoprotection, they were flash frozen in powdered dry ice and moved to a −80°C freezer for storage until sectioning. Brains were then sectioned coronally on a sliding microtome at 30 μm thickness into six series (so subsequent sections in a single series were 180 μm apart) and immediately mounted and allowed to air dry before being placed in a vacuum chamber with humidity sponges where they were left to dry fully for 24 hours. Only hippocampal sections (approximately −3.2 to - 5.2 from bregma) or prefrontal regions (approximately +3.7 to +1.4 from bregma) containing the areas of interest were sectioned and processed. Mounted sections were then placed in a sealed slide box and stored in a −80°C freezer until processing.

#### Tissue processing

All procedures were modified from the protocols used in Lee et al. [38] and Petrovich et al. [39]. Prior to tissue processing, a cRNA probe for *Arc* mRNA was constructed starting with a plasmid containing a full-length cDNA (~3.0 kbp) of the *Arc* transcript. To create the probe, the DNA was first cut by mixing the plasmid with a 10x digestion buffer (NEBuffer; Biolabs; Ipswich, MA, USA), a 10x EcoRI restriction enzyme (Biolabs), and purified nuclease free water (Ambion) before being incubated at 37°C for 2 hours. Proper cutting of the DNA was verified using electrophoresis, after which the DNA was purified overnight in ethanol. Following purification, the DNA pellet was spun out in a centrifuge, washed in EtOH, fully dried, and resuspended in a TE buffer. To verify that the DNA was properly linearized, calculate *Arc* concentration, and check that no contaminants were present, a sample of the DNA was tested via spectrophotometry (Nanodrop Lite; Thermo Scientific, Waltham, MA, USA). The Digoxigenin (DIG) labelled probe was transcribed by combining the linearized DNA with RNase free water (Ambion), a 10x transcription buffer (Ambion), RNAse block (Ambion), DIG RNA labelling mix (Roche Applied Science; Indianapolis, IN, USA), and a T7 RNA polymerase (Ambion) before incubating the solution at 37°C for 2 hours. Finally, the probe was diluted in nuclease free water and purified in a mini Quick-Spin column (Roche).

Once the cRNA probe had been constructed, slides containing tissue from the male rats were submerged for 40 minutes in a 4% PFA solution to increase tissue integrity throughout *in situ* processing. Tissue from female rats were processed without this PFA wash. Slides were then washed and incubated in a proteinase K (PK) buffer at 37°C before being treated with a 0.5% acetic anhydride/1.5% triethanolamine solution containing glacial acetic acid for permeabilization. Slides were then washed in a saline-sodium citrate (SSC) buffer before being dehydrated by submersion in ascending concentrations of ethanol and air dried. Finally, each slide was covered in 300 μl of a hybridization buffer containing yeast tRNA (Invitrogen; Carlsbad, CA, USA), salmon sperm DNA (Ambion), dithiothreitol (Sigma; St. Louis, MO, USA), and the cRNA probe. Each slide was cover slipped and temporarily sealed using a DPX mountant (Electron Microscopy Sciences; Hatfield, PA, USA) before being incubated in the hybridization solution for 20 hours at 60°C.

Once hybridization was complete, cover slips were carefully removed, and slides were incubated in a 4xSSC buffer mixed with sodium thiosulfate (ST) at 60°C for an hour before being treated with an ethylenediaminetetraacetic acid-based solution to inhibit RNAse activity at 37°C. Following this, slides were washed in descending concentration of SSC solution mixed with ST again at 60°C. Tissue was then washed in a detergent solution (Tween20; Sigma) before being stained with the PerkinElmer TSA Fluorescein system (NEL701001KT; PerkinElmer, Waltham, MA, USA). Slides were placed in a humid chamber and treated with blocking buffer followed by an anti-DIG-HRP conjugate for 2 hours. Slides were then briefly washed in the detergent solution before being returned to a dark humid chamber and coated with a solution containing fluroscein tyramide reagent (FITC) and allowed to sit for 30 minutes. Finally, slides were washed, allowed to air dry, and cover slipped with a mountant containing the nuclear stain 4’,6-diamidino-2-phenylinodole (DAPI) (Vectashield; Vector Lab, Burlingame, CA, USA). Slides were stored in the dark at −20°C until imaging.

#### Imaging

All imaging was completed using an Axio Scope A1 microscope (Zeiss; Thornwood, NY, USA). Regions of interest were identified via DAPI staining using a 10x objective with the assistance of the Paxinos and Watson brain atlas [40] and then imaged under a 40x objective (actual magnification ~900 X). Images were taken for both DAPI and FITC stains and later colorized and merged automatically using a custom macro in the ImageJ software with FIJI (NIH, Bethesda, MD). Due to tissue damage occurring over the course of *in situ* not all sections or areas of potential interest were viable. As such, images were not able to be z-stacked reliably and, instead, were taken on a single plane. The following regions were imaged and counted: the prelimbic cortex (+3.72 to +2.52 from bregma), the infralimbic cortex (+3.52 to +2.2 from bregma), the lateral (+3.72 to +3.2 from bregma) and ventral (+3.72 to +3.0 from bregma) orbitofrontal cortex, the CG1 region of the anterior cingulate cortex (+3.72 to +2.52 from bregma), and the CA3 region of the ventral hippocampus (−4.3 to −4.8 from bregma) (See Fig 3). Though the amygdalar nuclei were also of particular interest for their well-established role in fear learning, the aforementioned tissue damage tended to be particularly severe in this area. As such, we were not able to obtain a sample size large enough to include that region (a minimum of 6 viable images/region was required for a rat to be included in the statistical analysis of a given area).

**Fig 3.**
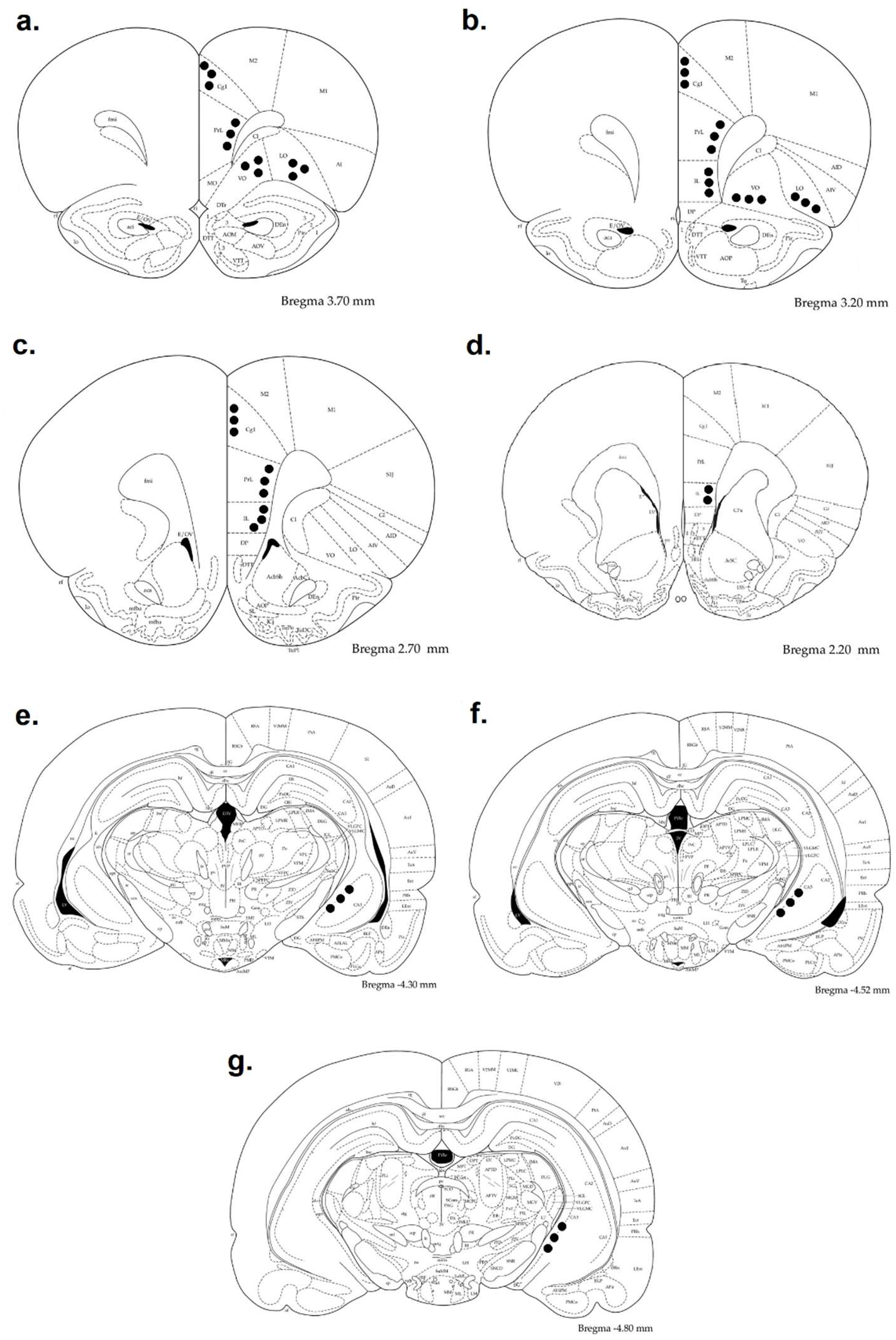
Representation of sampled areas. Images of coronal rat brain sections adapted from Paxinos and Watson (2006). The blacked-out circles indicate the approximate areas sampled from each plane for (a-d) the prelimbic, infralimbic, CG1 region of the anterior cingulate cortex, and the ventral and lateral orbitofrontal cortices, and (e-g) the CA3 region of the ventral

**Fig 4.**
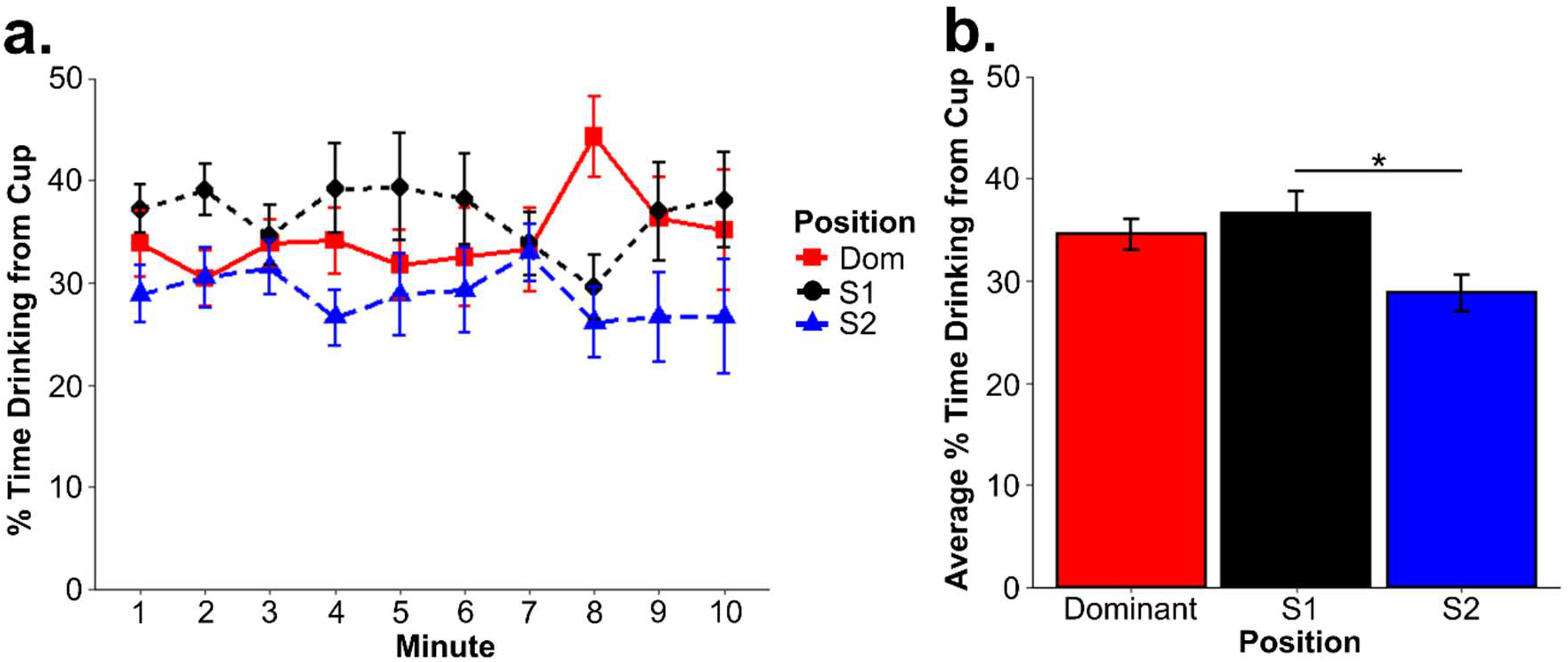
Milk dominance test results. The above figures show the average percent of total time that rats assigned a given rank spent monopolizing the milk cup (a) across the first ten minutes of the dominance test and (b) averaged across each minute by rank. *\*p < 0.05* rats and S1 rats (p = 0.713) (see Fig 4). Notably, these results are counter to earlier findings from our lab [13], which might be attributable to differences in the container used to hold the milk during testing as the lid of our container was slightly wider (4.45 cm Diameter vs. 3.75 cm diameter) making the milk more easily accessible.

Counts were completed region by region and all image files were assigned a random numerical code to blind the experimenter completing the counts from any details concerning the image at the time of counting. All cell counts were taken in ImageJ with the FIJI package and were made using the cell counting tool. Cells were counted for nuclear and cytoplasmic *Arc* mRNA expression separately and cells showing overlapping expression were counted as dual expressing. The final counts for nuclear *Arc* expressing and cytoplasmic *Arc* expressing cells included only those cells expressing in only that region (i.e., did not include dual expressing cells). Full counts for DAPI stained cells were taken and the percent of cells showing expression in each given area was calculated followed by the average percent of cells showing each type of activation in individual rats. To prevent the scores of rats with larger numbers of images from having a disproportionate effect on our statistics and to prevent an inflation of sample size only these averages were used in our final analysis.

## Results

All statistical analyses were completed using the R coding software. The full code is freely available to view at our data repository at https://dataverse.tdl.org/dataverse/MonfilsFearMemoryLab. Unless otherwise stated, the cutoff for a test to be considered statistically significant was set to p < 0.05.

### Behavioral Results

#### Dominance tests results

As only Cohort 2 rats were assigned conditions based on dominance rank, Cohort 1 males were not included in these analyses, resulting in data from nine triads (n = 27 rats) being included. To verify our dominance assignments, we ran a two-way ANOVA (Type 2) with the percent of total nape contacts in the cage received as the dependent variable and engaging rat rank and responding rat rank as independent variables. The interaction had to be tested separated using a one-way ANOVA. We found an overall effect of both engaging (F_(2,49)_ = 16.409, p < 0.0001) and responding (F_(2,49)_ = 19.490, p < 0.0001) rank and an interaction between the two (F_(5,48)_ = 9.84, p < 0.0001). A post-hoc Tukey HSD found that, as expected, dominant (p = 0.0082) and S1 (p = 0.043) were significantly more likely to engage than S2 rats, and dominants were more likely to be the responder when compared to both S1 (p = 0.0031) and S2 (p = 0.002) rats. S1 rats were also significantly more likely to contact the dominant rat than the S2 rat (p = 0.00031). We also examined the percent of times a rat responded to a nape contact with a counter, a behavior that has previously been found to be more likely in dominant rats [37]. Differences in likelihood of counter response was tested using a series of Kruskal-Wallace tests due to violations of ANOVA assumptions. We found that while there was no detected effect of engaging rank (H_2_ = 3.23, p = 0.199) there was a significant effect of responding rank (H_2_ = 7.92, p = 0.0191) and a post-hoc Dunns test with Holm’s p-adjustment found that dominant assigned rats did counter significantly more than rats assigned to the S1 condition (p = 0.0162) but not rats assigned to the S2 condition (p = 0.156). A mixed-effects ANOVA run to examine performance during the milk dominance assessment with percent of time monopolizing the milk cup as the dependent variable, assigned rank as the between-subjects variable, and minute of scoring as the within-subjects variable. We found a significant overall effect of rank (F_(2,24)_ = 4.83, p = 0.0172) but no effect of minute (F_(9,216)_ = 0, p > 0.99) and no interaction between the two (F_(18,216)_ = 1.189, p = 0.272). Post-hoc pairwise comparisons across the various ranks averaged across minute found that S1 ranks rats spent significantly more time monopolizing the milk cup than S2 rats (p = 0.0165) and dominant rats also trended in that direction (p = 0.09), but there was no significant difference between dominant

#### Fear conditioning and fear conditioning by-proxy

To ensure that Demonstrators had sufficiently acquired the CS-US association, their freezing on day 2 (during the FCbP observation period) was run through a one-way within-subjects ANOVA with timepoint (pre-CS, CS1, CS2, and CS3) as the within-subjects factor. We found a significant effect of cue (F_(3,87)_ = 77.96, p < 0.0001) and a post-hoc pairwise testing using a Bonferroni adjustment for multiple comparisons confirmed that freezing during the CS was significantly higher than at baseline (all p < 0.001) (see Fig 5a). A set of Kruskal-Wallace analyses were run on freezing on to the CS presentation on the terminal day (day 4) of the experiment as a nonparametric alternative to an ANOVA due to violations of ANOVA assumptions by the untransformed dependent variable. Kruskal-Wallace analyses were run on sex, experimental condition, and a factor containing all combinations of the two (to detect potential interactions) as independent variables. It found that while there was no overall effect of sex (H_1_ = 1.55, p = 0.2132) on its own, there was a significant effect of experimental condition (H_2_ = 35.1, p < 0.0001) and a significant effect of the combined factors (H5 = 37.38, p < 0.0001). Post-hoc Dunn’s tests using the Holm adjustment for multiple comparisons found that rats in the Demonstrator condition froze significantly more to the CS than both Observers (p < 0.0001) and Controls (p < 0.0001), but, surprisingly, Observers and Controls did not significantly differ in their freezing from each other (p = 0.814). Dunn’s testing on the combined sex and condition variable found that the overall effect detected via Kruskal-Wallace was driven entirely by the Demonstrator condition, i.e., no interaction effects were detected (see Fig 5b). Notably, our Demonstrators also displayed an unusually low percentage of freezing to this final CS (mean = 25.2) that we were unable to replicate using near identical behavioral procedures (see Supporting Information methods). We did, however, confirm that the Demonstrators’ freezing during the CS period was not just context based by using a Wilcoxon signs-rank test (due to violation of the assumption of normality because of a floor effect for pre-CS freezing) to compare freezing during the CS to their freezing prior to CS presentation (Z = 49, p = 0.0013). That Observer rats did not show higher freezing than Control rats during the final CS presentation, while somewhat concerning, is likely the result of our using only a single CS presentation. While past research in our lab has found that FCbP observer rats will freeze over controls on the first CS presentation of a long-term memory test [12], there were some methodological changes (pre-exposure of controls to the CS and rats being run during their dark cycle) that resulted in slight changes in behavior. This was confirmed in a follow-up experiment run under similar conditions where we ran a full three CS recall test (see Supporting Information text).

**Fig 5.**
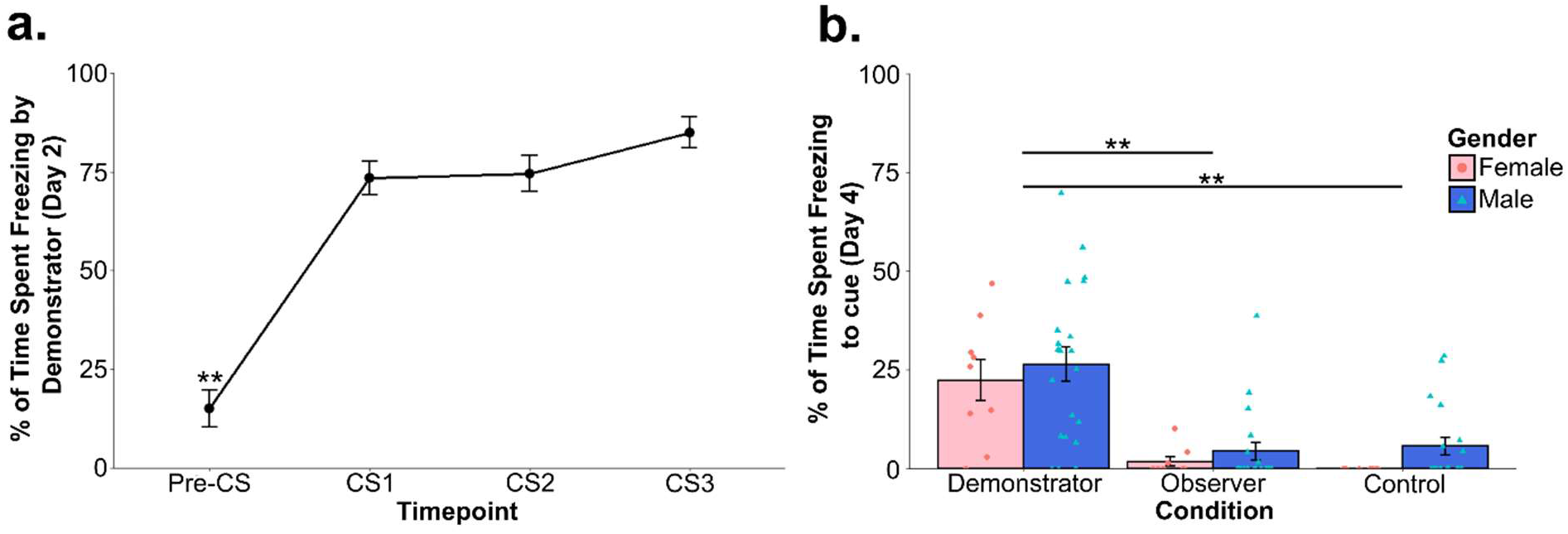
Fear conditioning and fear conditioning by-proxy behavioral results. The above figures show the average percent of total time that rats froze during or prior (Pre-CS) to the CS presentation for (a) Demonstrators on day 2, during FCbP interactions and (b) all rats to the single CS presentation on the terminal day of the experiment. *\*\*p < 0.005*

#### Choice test/food tasks

Choice test performance using either percent of time spent interacting the food cup containing diet Cin or percent of all eaten that was diet Cin was compared between Demonstrators and Observers using a two-sample t-test. We found no significant difference between the two groups on time spent at the diet Cin food cup (t_31_ = 0.97, p = 0.3404) or on the percent of total eaten that was diet Cin (t_31_ = 0.74, p = 0.4636). To determine whether this lack of an effect was due to both groups showing a preference for diet Cin, we ran a set of one-sample t-tests comparing the percent of total eaten that was diet Cin against the case in which rats showed no preference for either diet (μ = 50). We found that while both Demonstrator (t_16_ = 2.204, p = 0.04265) and Observer (t_16_ = 3.105, p = 0.0068) rats showed a significant preference for the diet Cin based on the percent eaten, neither Demonstrators (t_16_ = 0.476, p = 0.641) nor Observers (t_15_ = 1.885, p = 0.079) spent significantly more time interacting with the diet Cin food cup (see Fig 6a,b). The lack of difference between Observers and their Demonstrators can likely be explained by: (1) a slight innate preference for diet Cin over diet Co, as past research in our lab has found in Sprague-Dawleys [14], and (2) our decision to only use diet Cin as the demonstrated flavor in an attempt to decrease variance in the behavioral experience of our observers and (3) the brevity of the choice test compared to our standard design (10 minutes vs 1 hour). It is also worth noting that the Cohen’s d effect size for the Observer’s preference towards cinnamon (d = 0.75) is larger than the effect size calculated for Demonstrators (d = 0.53), though both fall into the category of medium effect sizes. Finally, to determine whether experimental condition influenced the total amount of food eaten, we ran a two-way ANOVA with total grams of food eaten during the choice as the dependent variable and experimental condition and sex as the independent variables. We found that while, as expected, there was a significant effect of sex (F_(1,82)_ = 35.66, p < 0.0001) (see Fig 6c) with females eating less than males, there was no significant effect of experimental condition (F_(2,81)_ = 0.334, p = 0.717) (see Fig 6d) and no interaction between the two (F_(2,81)_ = 1.02, p = 0.365).

**Fig 6.**
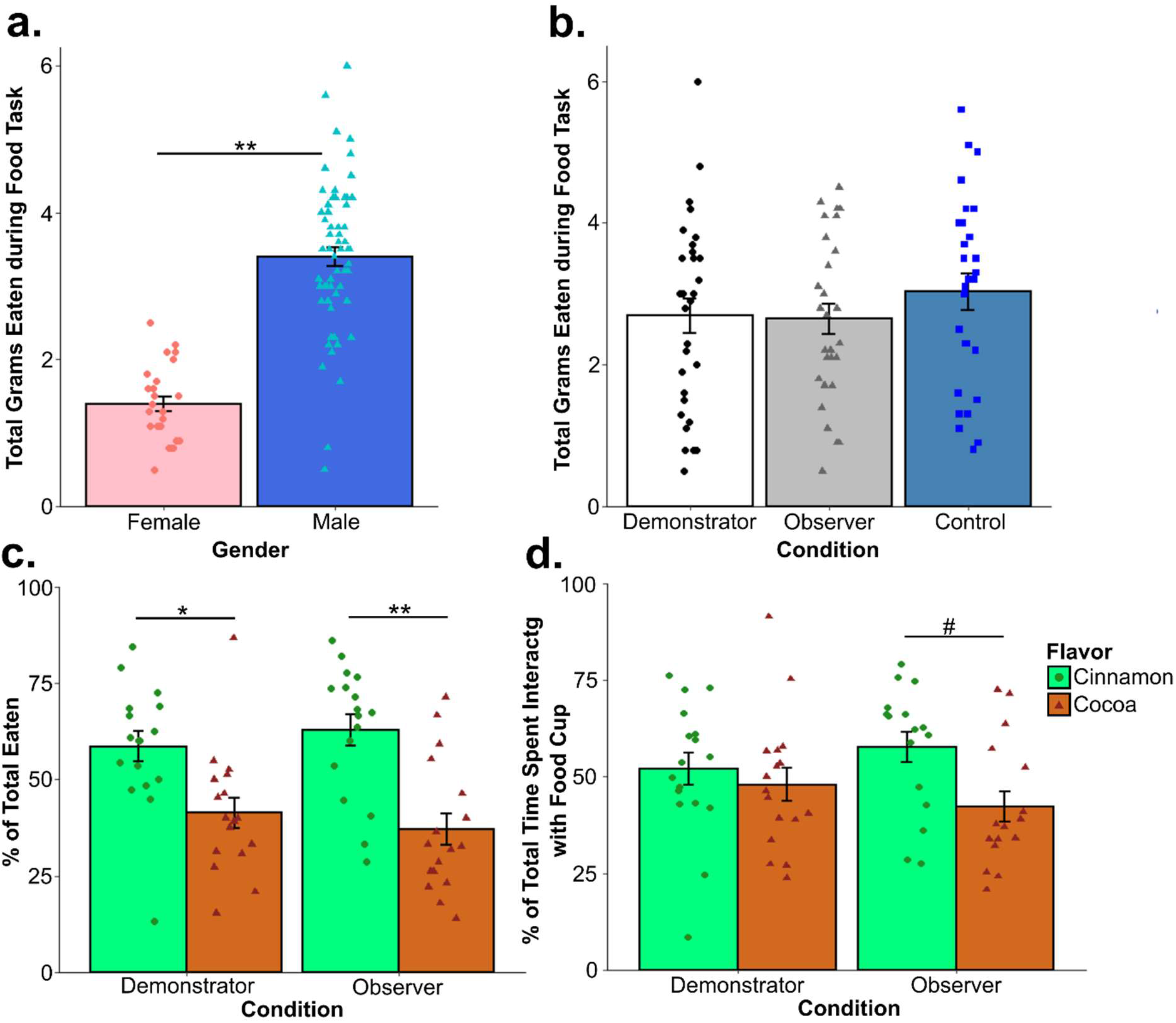
Day 4 food task behavioral results. For rats that went through the choice test on the final day of experimentation, we found that (a) while Observers and Demonstrators did not differ significantly from each other in the percent of diet Cin (the demonstrated flavor) eaten, they did both show a significant preference for the diet. However, (b) neither group spent significantly more time interacting with the food cup containing diet Cin. Examining the total amount eaten during the final food task for all rats we predictably found that (c) females overall ate significantly less than males but (d) experimental condition has no overall effect on the total amount eaten. *#p< 0.1, *p < 0.05, **p < 0.01*

### Arc Results

#### Arc Statistical analysis overview

All of our *Arc* results, unless otherwise mentioned, were tested for significance using a series of two-way ANOVAs (type 2) containing sex and condition as between subject variables (Sex and Condition) with an individual ANOVAs run for each area of expression (nucleus, cytoplasm, and dual). Similarly, a series of one-way ANOVAs were run with a combined variable containing the food task (diet Cin only or Choice test for Demonstrators and Observers; plain chow only for all Controls) for each area of expression. Sex was not included as a secondary variable as the relatively low number of female rats made sample sizes too small for certain condition/food task combinations. When ANOVA assumptions were violated, data were transformed using either a log(y+1) function or by taking the inverse square root. As these transforms did not always succeed in bringing ANOVAs in line with assumptions, Kruskal-Wallace tests were performed on datasets where transforms were not effective. Pairwise t-tests were performed for post-hoc analyses against a Bonferroni-corrected alpha value when ANOVAs indicated a significant effect of condition (α = 0.017) or a significant sex and condition interaction (α = 0.008; conditions tested against each other within each sex only) with between-group effect sizes calculated using Cohen’s d. To provide a better gauge of variability for our smaller group sizes, the MS_Error_ obtained from our ANOVA was used in the denominator of post-hoc t-tests. Effect sizes for ANOVAs were calculated using the standard partial η^2^ formula and for Kruskal-Wallace tests using the formula η^2^_H_ = (H - k + 1)/(n – k). For simplicity of data presentation, unless the addition of the food task grouping variable resulted in a significant effect or unless a significant contribution of sex was detected all data were presented graphically split up by area of expression and overall experimental condition only. Any rats that had fewer than 6 viable images counted in a given brain region were excluded from the analysis for that area.

Bivariate correlations were calculated for Observer and Demonstrator animals to assess potential relationships between behavioral measures and *Arc* cell counts for expression occurring at appropriate timepoints (e.g., cytoplasmic *Arc* counts for percent of cinnamon eaten). Pearson’s correlation coefficients were used in the event that no outliers in either dataset were detected with a Grubbs test; if outliers were detected, Spearman’s correlation coefficient was used instead. To gauge whether a relationship between social learning and *Arc* in dual expressing cells in Observers, an overall metric of social learning – referred to from here on out as the social learning metric (SLM) - was calculated take the mean of the z-score standardized scores for the percentage of total eaten that was the demonstrated food and the percentage of time spent in contact with the Demonstrator during the FCbP social learning phase during the CS presentation (males) or after the CS presentation (females). Notably, percent freezing to the cue on the final day was not used for Observer rats because our results and the results of our follow up experiment (see Supporting Information data) indicated that the conditions of our behavioral testing procedure resulted in some freezing behavior even in Control rats – at least in males - and, as such, might not be the best gauge of the strength of the socially acquired fear response. As such, given our past findings that interactions with the Demonstrator during or after the CS (depending on sex) highly predicted later freezing to the cue [12–14], interaction with the Demonstrator at the sex appropriate timepoint was tested for correlations against nuclear *Arc* activity rather than freezing to the cue on the final day for Observer rats. For Demonstrators, a similar metric was calculated based on standardized freezing to the cue on the final day and the percent of total eaten that was the familiar diet (Diet Cin) and checked against dual *Arc* activity. To correct for the multiple tests run on each behavioral dataset (6, for each brain region), the critical p-value for correlations was Bonferroni adjusted to 0.0083.

#### Arc Results

(An overview of statistical results for each area can be found in S1-S4 Tables)

No significant effect of condition and no interaction between sex and condition was detected in the vCA3, infralimbic cortex, anterior cingulate cortex, or the lateral and ventral orbitofrontal cortex (all p > 0.05) (see Fig 7). Additionally, none of the one-way ANOVAs found a significant effect of condition when rats were further separated based on the food task they were assigned in any of these areas or in the prelimbic cortex (all p > 0.1). An overall effect of sex was found in a number of regions including: nuclear *Arc* expression in the ventral orbitofrontal cortex (F_(1,64)_ = 4.851, p = 0.031; η^2^_partial_ = 0.07); dual expressing cells in the lateral orbitofrontal cortex (F_(1,57)_ = 6.18, p = 0.016, η^2^_partial_ = 0.094); nuclear (F_(1,69)_ = 35.470, p < 0.001, η^2^_partial_ = 0.325), cytoplasmic (F_(1,69)_ = 60.715, p < 0.0001, η^2^_partial_ = 0.463), and dual expressing (F_(1,69)_ = 9.84, p = 0.003, η^2^_partial_ = 0.124) cells in the vCA3 of the hippocampus; dual expressing cells in the CG1 region of the anterior cingulate cortex (F_(1,66)_ = 15.930, p < 0.001, η^2^_partial_ = 0.194); in nuclear expressing cells (F_(1,73)_ = 18.05, p < 0.001, η^2^_partial_ = 0.196) and dual expressing cells (F_(1,73)_ = 13.666, p < 0.001, η^2^_partial_ = 0.15) in the infralimbic cortex (see Fig 8); and in cytoplasmic expressing cells (H_1_ = 4.3, p = 0.038, η^2^_H_ = 0.045) and dual expressing cells (F_(1,70)_ = 18.11, p < 0.001, η^2^_partial_ = 0.203) in the prelimbic cortex (see Fig 9b,c). Female rats displayed higher *Arc* counts than males in areas other than the anterior cingulate, infralimbic, and prelimbic cortices, in which male counts were higher across all conditions. Notably, post-hoc analyses found no overall significant effect of condition within the *Arc* counts for across any of the tested regions or areas of cell expression (all p > 0.1). The two-way ANOVA examining nuclear expression in the prelimbic cortex found a significant interaction effect between sex and experimental condition (F_(2,70)_ = 3.96, p = 0.023, η^2^_partial_ = 0.102). Post-hoc testing found a significant difference between nuclear *Arc* expression in female Demonstrators and female Controls only (t_9.7_ = 3.9, p = 0.0032, d = 1.22) (see Fig 9a). Correlational analyses found a significant negative relationship between the SLM score of Observer rats and the percent of cells showing dual *Arc* expression in the ventral orbitofrontal cortex (t_10_ = −3.41, p = 0.0066, r = −0.73) (see Fig 10a). Follow up analyses confirmed that this relationship was not significant when looking at either the standardized measure of percent cinnamon eaten (t_10_ = −1.795, p = 0.103, r = −0.49) or the standardized measure of sex relevant contact during FCbP (t_10_ = −0.75, p = 0.47, r = −0.23) alone (see Fig 10b,c). All other correlational analyses were not significant beyond our Bonferroni corrected alpha value (all p > 0.01).

**Fig 7.**
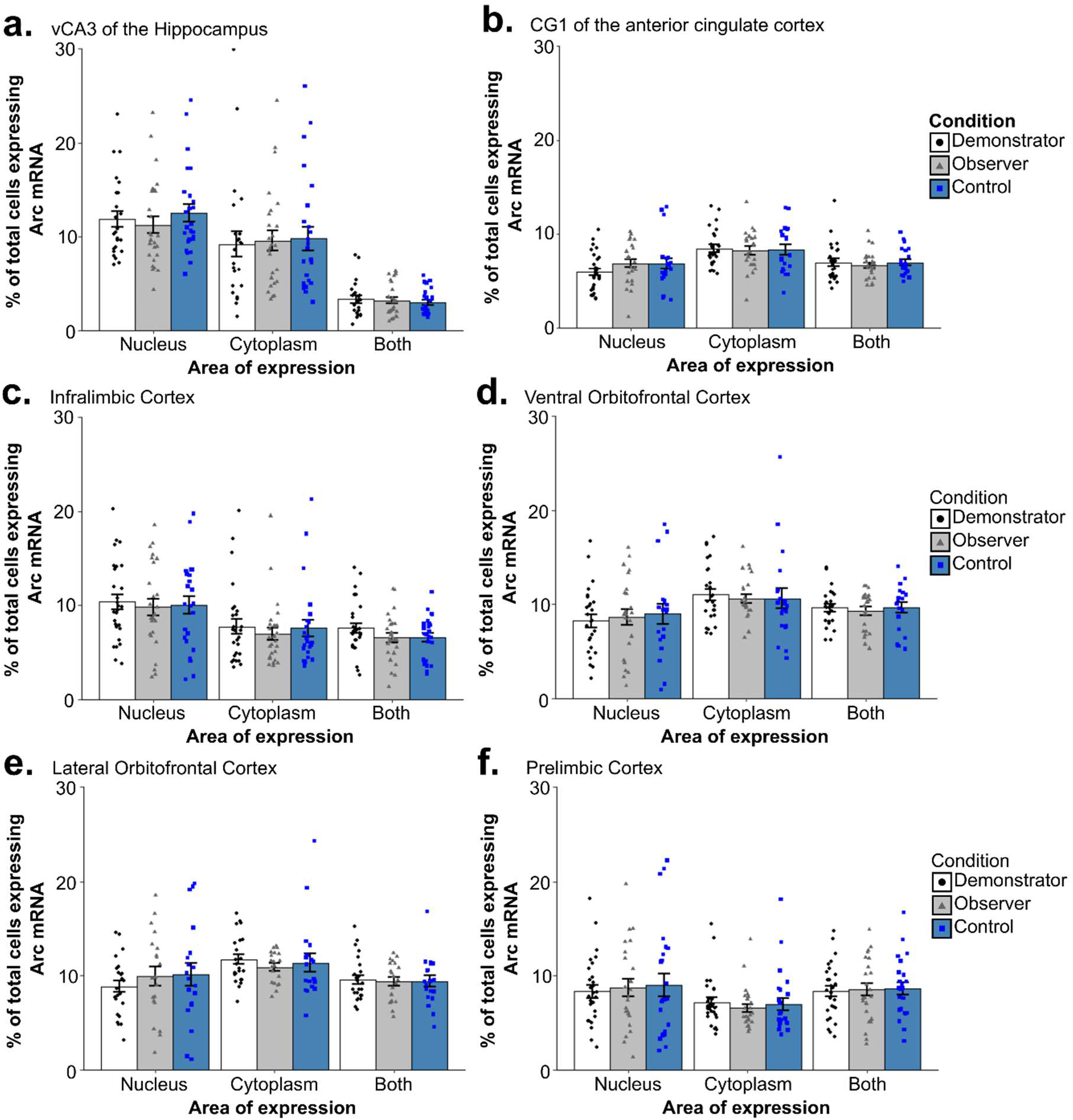
Arc counts across primary experimental condition. The above graphs show the percent of total DAPI stained cells that displayed *Arc* expression in the nucleus, cytoplasm, or in both area (dual) across the primary experimental conditions in (a) the vCA3 of the hippocampus, (b) the CG1 region of the anterior cingulate cortex, (c) the infralimbic cortex, (d) the ventral orbitofrontal cortex, (e) the lateral orbitofrontal cortex, and (f) the prelimbic cortex. Across all regions and areas of cell expression examined, no group differences were found between any of the conditions (all p > 0.1).

**Fig 8.**
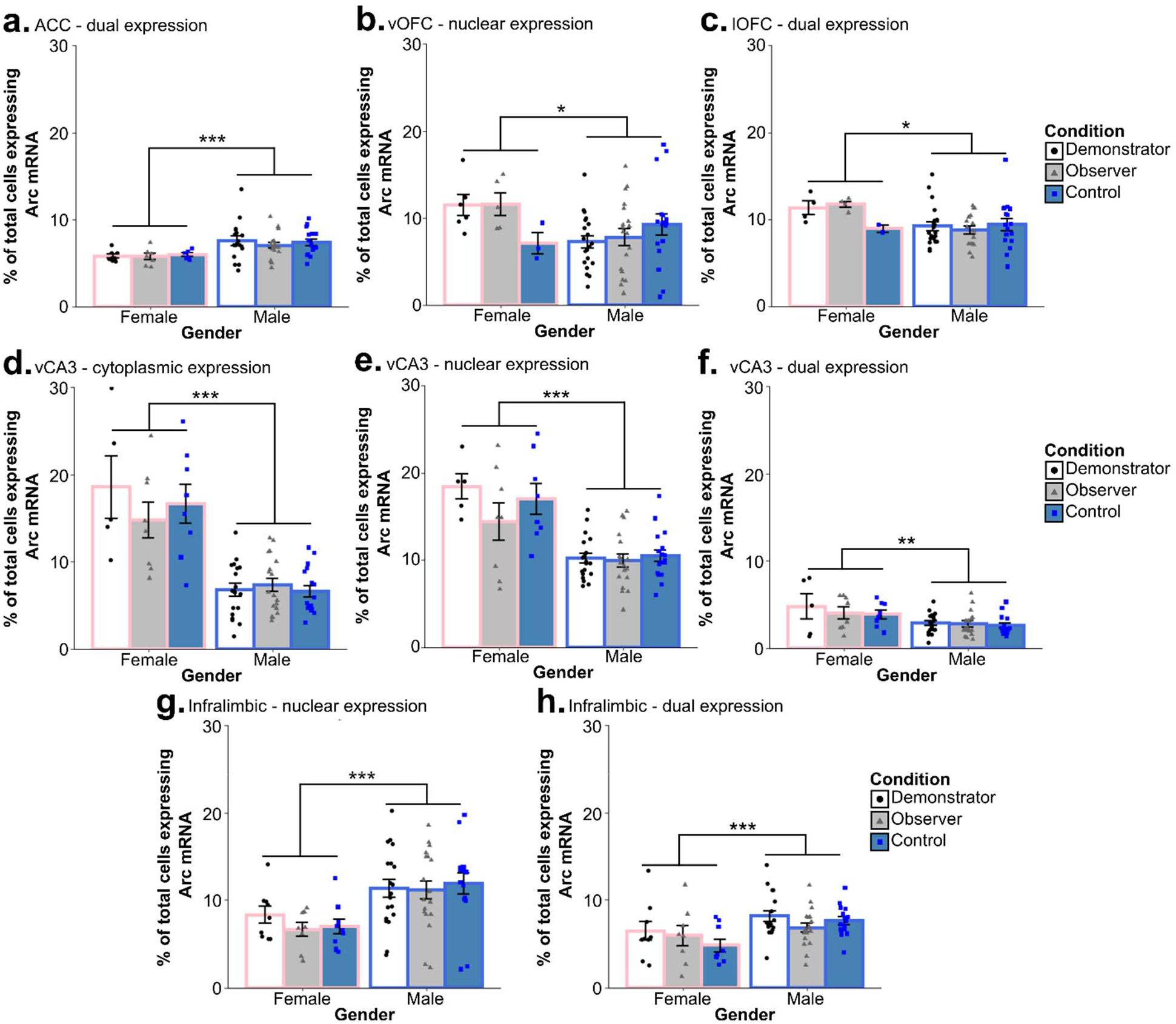
Differences in *Arc* expression between male and female rats. Significant differences in *Arc* expression were between male and female subjects when comparing (a) dual *Arc* expression in the CG1 region of the anterior cingulate cortex, (b) nuclear *Arc* expression in the ventral orbitofrontal cortex, (c) dual expression in the lateral orbitofrontal cortex, (d) cytoplasmic, (e) nuclear, and (f) dual *Arc* expression in the vCA3 of the hippocampus, and (g) nuclear and (h) dual *Arc* expression in the infralimbic cortex. +p < 0.1, *p < 0.05, **p < 0.01, ***p < 0.001

**Fig 9.**
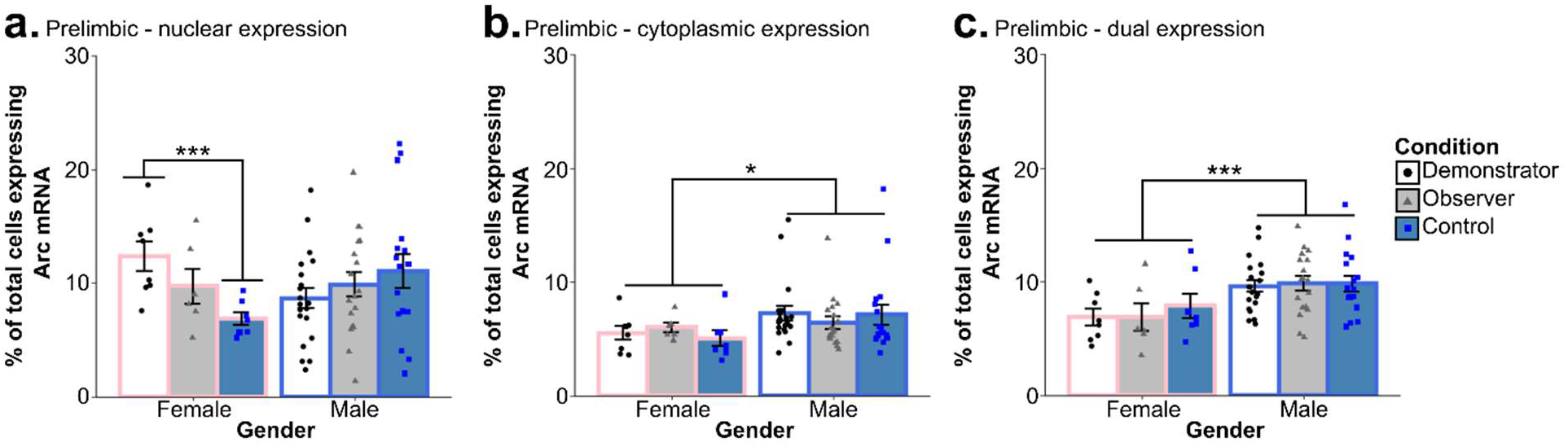
Differences in *Arc* expression between male and female rats across the prelimbic cortex. While initial ANOVA analysis found a significant sex and condition interaction in (a) the nuclear prelimbic counts, with female Demonstrators showing significantly more *Arc* expression than female Controls. Females did show lower overall (b) cytoplasmic and (c) dual *Arc* expression as compared to males in the prelimbic cortex, however. *\*p < 0.05, ***p < 0.001*

**Fig 10.**
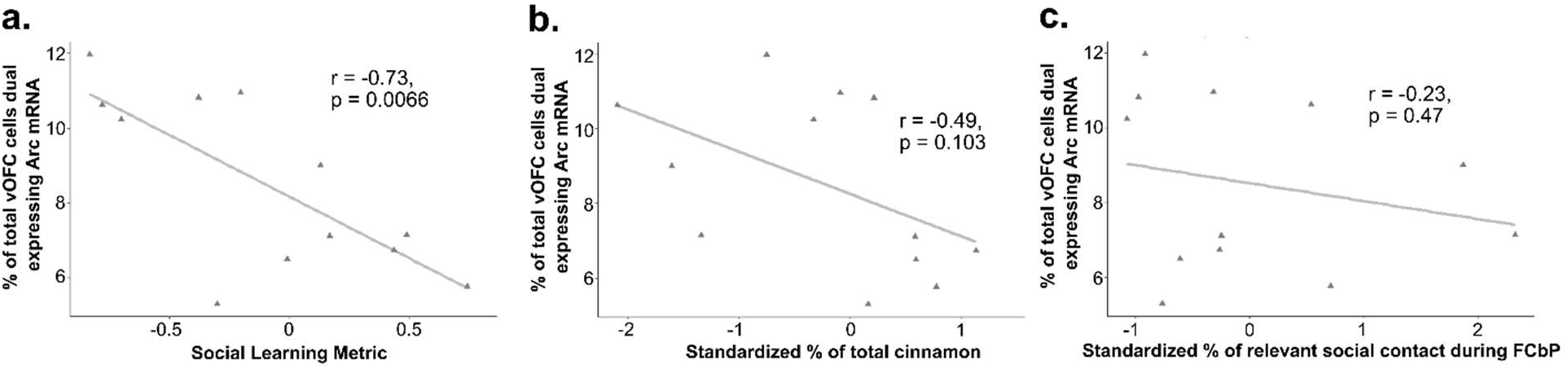
Relationship between social learning measures dual *Arc* expression in the vOFC. (a) A significant negative relationship was found between a social learning metric calculated by summing standardized measures of social acquisition of the STFP and socially acquired fear association in Observers and the percent of *Arc* dual-expressing cells in the ventral orbitofrontal cortex. This relationship was not significant when looking at either (b) the standardized measure of STFP or (c) the standardized measure sex relevant social contact during FCbP – used as a proxy for social fear learning - alone. Notably, both male and female animals were included in this dataset.

## Discussion

Contrary to our expectations, our results did not show any differences in *Arc* expression following long term memory recall based on whether the subject had acquired reward- and fear-based information by means of direct learning or social learning. Even more puzzlingly, Control rats that were put through analogous behavioral procedures prior to euthanasia but that had not been through any explicit fear- or reward-based training did not differ in *Arc* expression across the CG1 region of the ACC, the infralimbic cortex (IL), the vCA3 of the hippocampus, or the ventral or dorsal orbitofrontal cortex (OFC) when compared Demonstrators or Observers. Overall, the only differences in *Arc* expression that were detected were driven by subjects’ sex and showed no interaction with experimental condition. Though it is true that recall processes may not necessarily induce as many of the long-term changes in neural activity and connectivity that *Arc* is thought to be involved in [41] as learning procedures do, past research has found certain recall procedures to be sufficient to induce increased *Arc* activity [42,43]. As such, the lack of an effect across conditions that we see cannot be attributed only to our choice to examine learning at the recall timepoint. In the following sections, we will first examine our overall findings in the context of past research into the brain mechanisms underlying recall processes in the STFP paradigm, fear-conditioning and observational fear-conditioning procedures, and our findings in the ventral orbitofrontal cortex in the context of past research. We will then cover our findings – and, importantly, the limitations around our ability to interpret these findings – on sex effects on *Arc* expression.

### *Arc* in the recall of a socially transmitted food preference

Past research examining expression of the IEG c-Fos has found that a number of the areas we examined, specifically the orbitofrontal cortex, vCA3, infralimbic cortex, and the prelimbic cortex [22,23] show activation at the 48 hour recall timepoint for a socially transmitted food preference. It is also notable that these results from Smith et al. [23] were obtained using the same STFP control paradigm as was used in this study, indicating that though STFP recall induced activity in these regions may have been detectable with c-Fos, this may not be the case at this timepoint when examining *Arc*. This interpretation is backed up by the findings of Pilarzyk et al. [43], who examined *Arc* mRNA activity following STFP recall in Pde11a knockout mice, which displayed impaired recent STFP and enhanced remote STFP compared to Pde11a wild-type controls. They found that both animals showed increases in *Arc* expression over home-cage controls at this timepoint in the ventral and dorsal CA1, the ventral and dorsal subiculum, and in the CG1 and CG2 of the ACC. Moreover, while Pde11a knockout mice showed decreased *Arc* expression following a recall procedure for a recently acquired (24 hours post) STFP memory when compared to Pde11a wild-types in the vCA1, no difference between the two genetic lines was evident in any of the other regions examined. At a more remote recall timepoint (7 days post), knockout animals showed higher *Arc* activity post-recall in the CG1 and CG2 of the ACC but not in the vCA1 as compared to the wildtype controls, with home cage animals showing no baseline difference regardless of genetic line. Given that these differences in ACC *Arc* activity were not observed during early recall and the differences in vCA1 *Arc* activity was not seen during remote recall, it is reasonable to assume that this *Arc* activity was specific to both the experience of STFP recall and the recall timepoint. These findings are particularly interesting in light of prior research examining c-Fos activity in the vCA1 and the ACC at the respective timepoints at which enhanced *Arc* activity was seen in these animals, as past studies have found no differences in c-Fos activity in these areas when recall was induced on the exact same timeframe [22,23]. With this in mind, it is perhaps unsurprising that we also observed no recall induced changes in *Arc* expression in the various regions we examined despite their consistently being shown to be active using c-Fos as an activity marker. Exactly what the implications of this are - outside of the obvious conclusion that not all IEGs are equal – is hard to say when working with mostly null findings. That said, the high sensitivity of cellular compartment analysis of temporal activity by fluorescence *in situ* hybridization (catFISH) and our large group sizes for the primary behavioral conditions (Demonstrator, Observer, and Control) does lend validity to the non-significance of our findings. One caveat to our design that future experimenters might want to consider is the possibility that Demonstrators may also have acquired a STFP simply through exposure to the scent of the consumed food on their own breath and carbon disulfide from the nasal cavity of the Observer with whom they were interacting.

### *Arc* in the recall of direct and socially acquired fear associations

Our ability to interpret our findings regarding our rats undergoing recall of fear acquired via direct learning is significantly aided by how well-characterized the system underlying fear learning and recall is. A number of the areas we examined are well established as being involved in fear or extinction learning (the latter of which we would assume to be initiated in Demonstrators, as they had undergone non-reinforced CS presentation during FCbP) specifically the ACC, the prelimbic cortex, and the infralimbic cortex [30–32,44]. Though a much smaller pool of research is available regarding the neural mechanisms of social fear, the proposed models of social fear learning posit a system similar to that underlying recall of directly acquired fear associations also underlies the fear learning and recall processes for social fear learning [28]. Our findings indicate no overall role of the ACC, prelimbic cortex (PL), or IL in either recall of a socially acquired fear association or a directly acquired fear association (though see also discussion of sex differences in PL activity below). However, as covered in the previous section, this likely just indicates that *Arc* does not serve as a reliable indicator of activity in this case. Examination of these areas post-fear acquisition would likely tell a different story. Though explicit research in *Arc* activity following fear recall is limited, there is some past research to draw from. Chia & Otto [42] found that when Arc protein expression was examined following the presentation of a CS that a rat had acquired a fear association for via trace fear conditioning (i.e., fear conditioning with a delay between CS termination and shock delivery) rats were found to have significantly higher Arc expression in both the dorsal and ventral hippocampus when compared to unconditioned controls that were exposed to the chamber but not the CS. Notably, Arc was quantified by Western Blot analysis of the homogenized ventral and dorsal HPC in this experiment, so precise localization of HPC activity was not available. These findings likely indicate that, like in STFP, *Arc* transcription might be induced in certain areas of the hippocampus at the 48-hour recall timepoint for a cued fear memory.

### Potential Role of the Ventral Orbitofrontal Cortex in Recall of Socially Acquired Information

In a landmark study, Lesburguères et al. [24] were able to demonstrate that while dorsal hippocampal (dHPC) activity was necessary for acquisition and short-term recall of an acquired STFP, the STFP memory was eventually offloaded to the OFC for long-term storage. Additionally, Lesburguères et al. were able to demonstrate that that tagging of neurons in the orbitofrontal cortex during STFP acquisition is necessary for long-term storage of socially transmitted food preferences and that interference with the OFC activity following acquisition impairs remote memory recall (30 days post-acquisition) (though see also [45]). These findings would suggest ongoing communication between the dHPC and the OFC in the first days or weeks post-STFP acquisition and, furthermore, would suggest ongoing reorganization of the OFC at this timepoint to accommodate the long-term storage of the STFP memory. While the lack of overall differences in ventral or lateral OFC *Arc* expression between Demonstrators, Controls, and Observers in this study would challenge that interpretation somewhat, we did detect a significant negative correlation between our combined measure of overall social learning performance and dual-*Arc* expressing cells in the vOFC. Furthermore, this correlation was not observed between a similar metric formed for Demonstrators based on their choice test performance and their freezing to the cue. As reliance on socially acquired information can be thought of as making the choice between potentially unreliable social information and the potential dangers of learning through direct experience, it is possible that this apparent inhibitory role of the vOFC on expression of socially acquired information might be connected to the OFC’s broader role in value-based decision making [46–49].

### Sex Differences in *Arc* Transcription

Prior to this discussion, it should be stated that our ability to interpret our sex-related results is hindered for a number of statistical and methodological reasons. First, our occasionally low sample size for females, with group size for sex/condition combinations ranging from n = 2 to n = 9 following removal of rats without enough viable sections (though notably an n < 5 was only present for female Controls in the vOFC and lOFC and female Demonstrators and Observers in the lOFC). Additionally, our lack of entirely undisturbed controls means that we have no way to determine whether these sex differences are the result of baseline or task-specific differences in *Arc* mRNA production. Finally, because the pre-in *situ* PFA wash was not introduced until all female sections had been processed, it is possible that this difference in tissues processing might have affected the overall stain. That said, if this were the case, we might expect to see a broader and more consistent effect of sex across regions and types of *Arc* expression (nuclear, cytoplasmic, and dual). As it is, 18 regions/cellular areas of *Arc* expression combinations are examined and only 10 display a significant overall effect of sex. Furthermore, this effect is not uniform in its direction, with males displaying greater overall *Arc* expression in 5 cases and females displaying greater expression in the other 5. Regardless, we feel that our findings here should serve only to inform possible future research into sex differences in *Arc* expression. As it is, the limitation of the current study would make drawing definitive conclusions regarding sex effects on *Arc* expression inappropriate. This should be kept in mind in reading the following discussion.

Although there has been little investigation into sex differences in *Arc* expression, there are some findings indicating that female rats may show higher levels of *Arc* expression in certain regions of the dorsal hippocampus following repeated exposure to a relatively enriched environment [50], though a trend in the opposite direction has also been observed in animals tested without prior behavioral intervention [50]. Our findings may indicate that sex differences in *Arc* transcription may be present following certain general behavioral tasks or experiences. In the CG1 region of the ACC we found that males, overall, had more cells active at both timepoints, possibly due to higher baseline *Arc* transcription in the ACC of males or increased transcription following context changes/re-exposure (home cage → STFP testing room → conditioning chamber) as there is some evidence – though limited – for a role of the ACC in long-term recall of contextual memories [51]. Male rats also displayed higher nuclear and dual *Arc* counts in the infralimbic (IL) cortex. It is possible that the higher IL *Arc* counts in males might be explained by the role of the infralimbic cortex in extinction and fear inhibition [31,52,53] and the well documented impairments in the inhibition and extinction of learned fear in females [54–56]. If this is the case, however, it does raise the question of why no overall differences were observed between our Control, Observer, and Demonstrator animals if *Arc* expression was being triggered by CS-elicited infralimbic activity.

Females showed higher levels of *Arc* expression for all counts in the vCA3. The difference in nuclear counts could potentially have been the result of greater activation following exposure to the CS or re-exposure to the conditioning chamber in females, while the higher levels of cytoplasmic *Arc* expression in the vCA3 following the food task may indicate a sex differences in the role of *Arc* in the vCA3 either the recognition of “familiar” food (even for Observers the scent would be familiar due to their prior interaction with the Demonstrator) or reward/general consummatory processes. That females also showed significantly higher dual labelling in the vCA3 – though this effect was small – might also indicate generalized increases in vCA3 *Arc* transcription in females. Female rats also displayed higher nuclear *Arc* transcription in the ventral OFC and higher dual levels of *Arc* mRNA in the lateral OFC, though these results are more difficult to interpret due to the low number of female Control rats whose brain tissue was intact enough to take OFC counts (n = 2 and 3 for the lateral and ventral OFC, respectively). Data from the Control rats we do have indicate a possible sex mediated increase in OFC *Arc* mRNA production, but it is just as possible that this effect would not persist with a higher n. It is notable that some past research has indicated structural differences in the OFC and functional differences in OFC-mediated behaviors between female and male rodents [57–59].

Possibly our most interesting sex differences in *Arc* mRNA were detected in the prelimbic cortex. In the prelimbic cortex (PL), males showed overall higher numbers of cells expressing *Arc* in the cytoplasm and in both the cytoplasm and nucleus (dual expressing) than females. While no within-sex differences across condition assignments were detected for these counts, we did find a significant sex/condition interaction in our nuclear prelimbic counts. Specifically, it appears that while male Demonstrators and Observers did not show increases in *Arc* transcription over Controls at the fear-recall timepoint, female Demonstrators showed significantly higher nuclear *Arc* transcriptions than Controls while female Observers fell in the middle between the two. This sex-effect may be driven by the aforementioned deficits observed in learned fear inhibition and extinction that are observed in females [54–56], as past research has suggested that the PL is critically involved in stimulating fear behavior [52,60,61], essentially serving an opposing role to the IL. Furthermore, a number of studies have implicated differences in PL signaling and structure as potential driving factors for these sex-specific impairments in fear-inhibition and extinction [62–65]. While we found no significant difference in female and male freezing behavior to the cue, the upregulation of *Arc* mRNA in response to a non-reinforced fear associated CS in specifically female Demonstrators may be indicative of differential neural restructuring in the PL that could ultimately lead to sex differences in fear expression.

### Conclusions

While the findings of this study did not broaden our understanding of the brain mechanisms involved in the retrieval of socially acquired memories as much as we had hoped, our results do provide some potential insights on sex differences in *Arc* expression as well as the role (or lack thereof) of *Arc* in long-term memory recall. Our findings suggest that - at least in the prefrontal cortex and vCA3 – the induction of brain activity through recall of socially acquired information does not appear to be sufficient to cause increases in *Arc* expression over those caused by the testing procedure alone. However, the validity of this takeaway is certainly brought into question by the inconclusive results of our behavioral tests, which might suggest that poor retainment of the socially acquired information was at fault for this lack of effect. We theorize that this may be because minimal neural restructuring is triggered when recall occurs prior to systems consolidation. Further research into the role of the Arc protein in social learning recall processes is still warranted given that our behavioral results do not demonstrate social learning in Observer rats as definitively as we would have hoped. Future research examining overlap in the neural mechanisms governing different forms of social learning might also benefit from the inclusion of animals undergoing acquisition procedures and animals undergoing remote recall procedures, as these timepoints may be more likely to induce plasticity changes and thus changes in *Arc* expression. Though the short timeframe of *Arc* expression in and around the cell body may make this methodologically difficult to achieve, rapid acquisition of a STFP might be achieved by using multiple demonstrators at once.

## Declarations

### Author’s Contributions

LAA designed the experiment, gathered and analyzed data, processed and imaged tissue samples, and drafted the manuscript. ENH and JD assisted with tissue sample processing. VN assisted with the experiments documented in Supporting Information. MHM and HJL designed the experiment, gathered data, provided guidance and training for tissue processing and imaging, and approved the final version of the manuscript.

### Availability of data and material

Raw data files are available in The Monfils Lab repository, housed in the Texas Data Repository in Dataverse (https://dataverse.tdl.org/dataverse/MonfilsFearMemoryLab). All other materials are available by request to the authors.

### Code availability

Code is available alongside data and materials in the Monfils Lab repository (see above).

