## Supplementary Materials for "Patterns of Arc mRNA expression in the rat brain following dual recall of fear- and reward-based socially acquired information"

**Supplementary Methods and Results** p2-p7

**Supplementary Figures and Tables** p8-p15

**Contact Information:**

Marie-H. Monfils

The University of Texas at Austin

Department of Psychology

108 E. Dean Keeton Stop A8000

Austin, TX 78712-1043

### Supplementary Methods and Results

#### Fear Conditioning by-Proxy in Dark Cycle with Control Cue Exposure

Past behavioral experiments run in our lab using the fear conditioning by-proxy (FCbP) behavioral paradigm had animals run during their light cycle for all phases of the procedure and did not involve control animals receiving CS exposure prior to long-term memory testing (Bruchey, Jones, & Monfils, 2010; Jones & Monfils, 2014; Jones & Monfils, 2016; Agee, Jones, & Monfils, 2019). In order to determine whether the lack of difference in freezing between our control and observer animals on the final day of experimentation in the main experiment was due to the methodological changes that we had made in the fear conditioning by-proxy (FCbP) for the primary experiment, a follow up experiment partially replicating our FCbP procedures with a full long-term memory (LTM) test was run.

*Subjects and scoring*

Subjects were 24 male Sprague-Dawley rats (n = 8/condition) that had been housed in their triad for a month and had previously been run through a novel object experiment that took place in the conditioning chambers. Therefore, all rats were familiar with the conditioning chambers but had only ever interacted with a novel object within the chamber. One rat from each triad was randomly assigned to the control, observer, and demonstrator condition. Following the experimental procedure, videos of the long-term memory test were scored for freezing to just prior to the first cue (Pre-CS freezing) and freezing to all three subsequent cues using the same scoring method as described in the main paper.

*Methodology*

The FCbP procedure was run exactly as described in the main text with the following exceptions: (1) all animals were given three exposures to the conditioned stimulus (CS) during the long-term memory test on the final day, (2) control animals were run through cue exposure on the same day as FCbP acquisition took place for observer rats, and (3) the long-term memory test took place 24 hours following FCbP training/cue exposure rather than 48 hours after FCbP training/cue exposure, and (4) demonstrators and observers did not go through the social transmission of food preference (STFP) procedure after the FCbP interaction (see *Figure S1*). Notably, to prevent the behavior of observer and demonstrators (specifically, potential olfactory or auditory cues) from influencing controls during cue exposure, all control animals were run prior to observers and demonstrators being placed in the chamber.

*Results*

A two-way mixed ANOVA was run on the percent of time freezing during or just prior to CS presentation with cue period (pre-CS, CS1, CS2, and CS3) as the within-subjects variable and experimental condition as the between-subjects variable. A significant overall effect of condition ($F_{(2,21)} = 7.588$, $p = 0.0033$) and cue ($F_{(3,63)} = 10.68$, $p < 0.0001$) but no interaction between the two ($F_{(6,63)} = 0.73$, $p = 0.62$). Post-hoc Dunns tests with Holm's adjusted p-values for multiple comparisons found that demonstrators displayed significantly higher freezing than observers ($p = 0.0018$) and controls ($p < 0.00001$), but observers did not freeze significantly more than controls ($p = 0.11$). They also confirmed that the only significant differences in freezing between cue periods was between the Pre-CS period and the CS1 ($p = 0.014$) and CS2 ($p = 0.00563$) periods, though the difference between pre-CS and CS3 was only nearing significance after correction ($p = 0.078$) (see *Figure S2a*). A one-way ANOVA was also run on the percent freezing to cue averaged across the three CS presentations with condition as the between-subjects variable. While the ANOVA found an overall effect of experimental condition ($F_{(2,21)} = 16.95$, $p < 0.0001$), a post-hoc Tukey HSD again found that this difference was significant between demonstrators and observers ($p = 0.0009$) and demonstrators and controls ($p < 0.0001$) but not between controls and observers ($p = 0.428$) (see *Figure S2b*). These results back up our interpretation of the lack of difference in freezing between our observers and controls as likely being the result of our methodological changes to the FCbP paradigm.

**Effect of STFP and FCbP on freezing response in directly conditioned animals**

To determine whether the unusually low freezing that was documented in our demonstrator animals at the final long-term fear memory test was: (1) replicable and (2) the result of social component of the behavioral procedures that demonstrators had undergone during the second day of the experiment, a new set of rats were run through iterations of the behavioral procedure with varying amounts of social interaction.

*Subjects*

Subjects were 48 male Sprague-Dawley rats that were ~6 weeks of age at arrival. Rats were housed in triads immediately after arrival. Testing did not start until at least a month after arrival to ensure that the rats in each triad had sufficient time to form social bonds. Rats were left undisturbed until four days prior to the start of behavioral testing, at which point they were food restricted and habituated to handling procedures/testing areas as described in the main text. 16 rats served as familiar social stimulation only, resulting in an n = 8 group size for each of the four experimental conditions.

*Methodology*

(See *Figure S3* for a graphical overview of the behavioral design)

Rats in each triad were assigned to one of three conditions: (1) Demonstrator 1 (Dem1), (2) Demonstrator 2 (Dem2) or (3) Observer. As we were only interested in the behavior of the demonstrator (i.e., the directly fear conditioned rat) animals in this case, observer rats served purely as a familiar animal to provide the social component of the behavioral paradigm. On day 1 of the experiment, both demonstrators went through fear conditioning exactly as described in the main text. On day 2, demonstrators were re-exposed to the CS 3 times either in the presence of their paired observer or alone before either being transported to single housing or being given access to cinnamon flavored powdered chow for 1 hr, after which they were allowed to interact with their triad's observer for 30 minutes. This results in there being four possible experimental conditions: (1) Recall with cage mate and STFP (RC+STFP), (2) Recall with cage mate only (RC), (3) Recall alone only (RA), and (4) Recall alone and STFP (RA+STFP). The recall procedure was the same protocol as what was used for the FCbP in the primary experiment. As Dem1 and Dem2 were run at the same time, in triads in which one demonstrator was assigned to the RC+STFP condition the other demonstrator would always be assigned to the RA condition. Similarly, in cages in which one demonstrator was assigned to the RC condition, the other was always assigned to the RA+STFP condition. Following day 2 behavioral procedures, observer animals were euthanized and demonstrators were moved to single housing.

48 hours later, on the final day of behavioral procedures, all demonstrators were given access to cinnamon and cocoa chow and allowed 10 minutes to consume as much of either as they pleased. After this, both demonstrators were returned to their home cage for 10 minutes before being returned to the conditioning chamber and allowed to habituate for 5 minutes before being given three 20 second CS presentations. Three CSs were given rather than the single presentation in the primary experiment because we deemed it more important to be able to thoroughly investigate their behavioral response than to ensure clean *Arc* expression. Demonstrators were, however, perfused as described in the main text and brains were removed, cryoprotected, and flash-frozen in case the behavioral results warranted tissue analysis.

*Results*

A two-way mixed ANOVA was run on the percent freezing to the CS on recall day 2 with cue period (pre-CS, CS1, CS2, CS3) as the within-subjects variable and experimental condition as the between-subjects variable. A significant effect of cue period ($F_{(3,84)} = 17.49$, $p < 0.0001$), but not experimental condition ($F_{(3,28)} = 0.78$, $p = 0.52$), or any interaction between the two ($F_{(6,56)} = 1.14$, $p = 0.35$) was detected. A post-hoc Dunns test with Holm's correction confirmed the effect of cue was driven by freezing at the pre-CS period ($p < 0.001$ vs freezing during all CSs) (see *Figure S4a*). A set of planned pairwise comparisons (two-way t-tests with Holms adjusted p-values) were run to check for differences in freezing to CS1 only between the RC+STFP condition and each other condition. No significant differences were detected between rats in the RC+STFP condition and rats in any of the other conditions (all $p > 0.1$). Finally, a two-way factorial ANOVA run on the average percent freezing during each CS presentations found no significant effect of recall condition ($F_{(1,28)} = 0.06$, $p = 0.81$) or STFP condition ($F_{(1,28)} = 0.96$, $p = 0.34$) and no interaction between the two ($F_{(1,28)} = 0.46$, $p = 0.51$) (see *Figure S4b*). While the freezing to CS1 by the rats that were assigned to the behavioral condition that was most similar what the demonstrators had undergone in the main experiment (RC+STFP) (n = 8, mean = 43.2, SD = 24.0) was higher than the freezing observed in our demonstrators at the final long-term memory test of our primary experiment (n = 30, mean = 25.2, SD = 18.5), this difference did not quite reach significance when the scores of the demonstrators were compared against each other with a two-sample t-test ($t_{9.3} = -1.97$, $p = 0.079$). It is possible, if unlikely, that adding more animals to the RC+STFP condition would have resulted in their CS1 freezing scores regressing towards the mean of the freezing of our demonstrators in the main experiment. In interpreting these results, it is also important to note that the social relationship between the observers and demonstrators in this experiment were quite different from the relationship between observers and demonstrators in the original experiment (unrelated 1-month cage mates vs. siblings housed since weaning) and this may explain the relatively lower freezing observed in the demonstrators in our primary experiment.

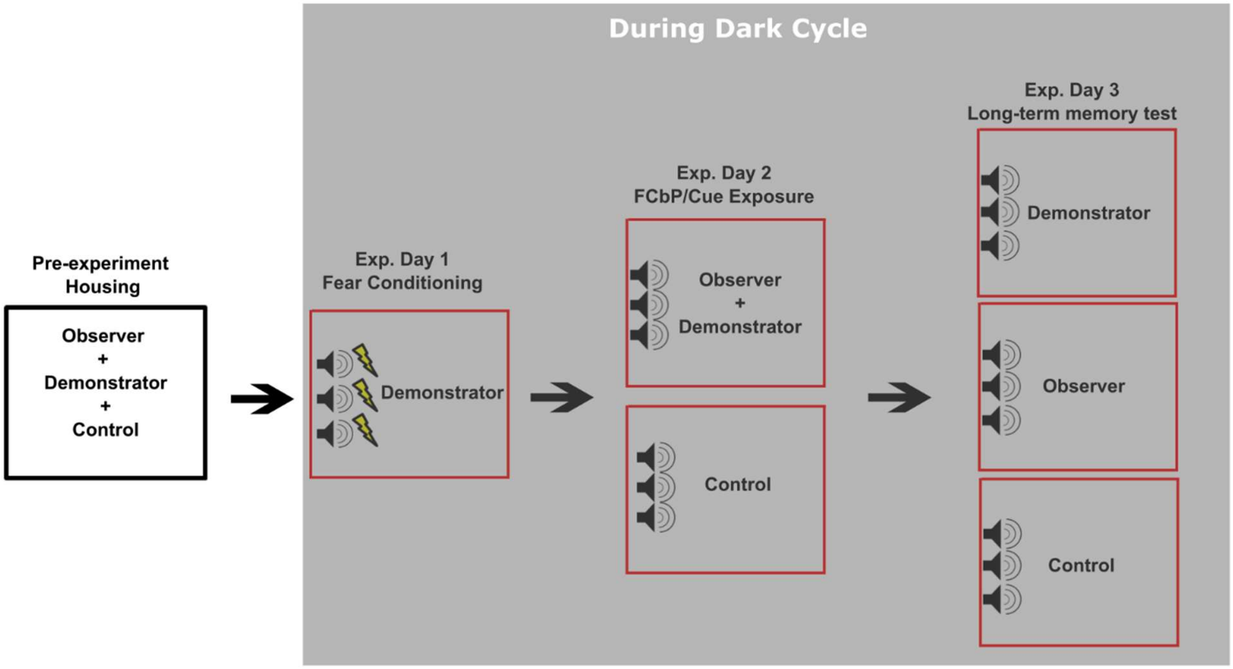


***Figure S1.*** *Dark cycle fear conditioning by-proxy with control cue exposure.* The above figure outlines that procedure for our follow-up experiment examining long-term freezing response to an auditory CS in rats that had undergone FCbP during their dark cycle when compared to control animals that had received cue-exposure alone.

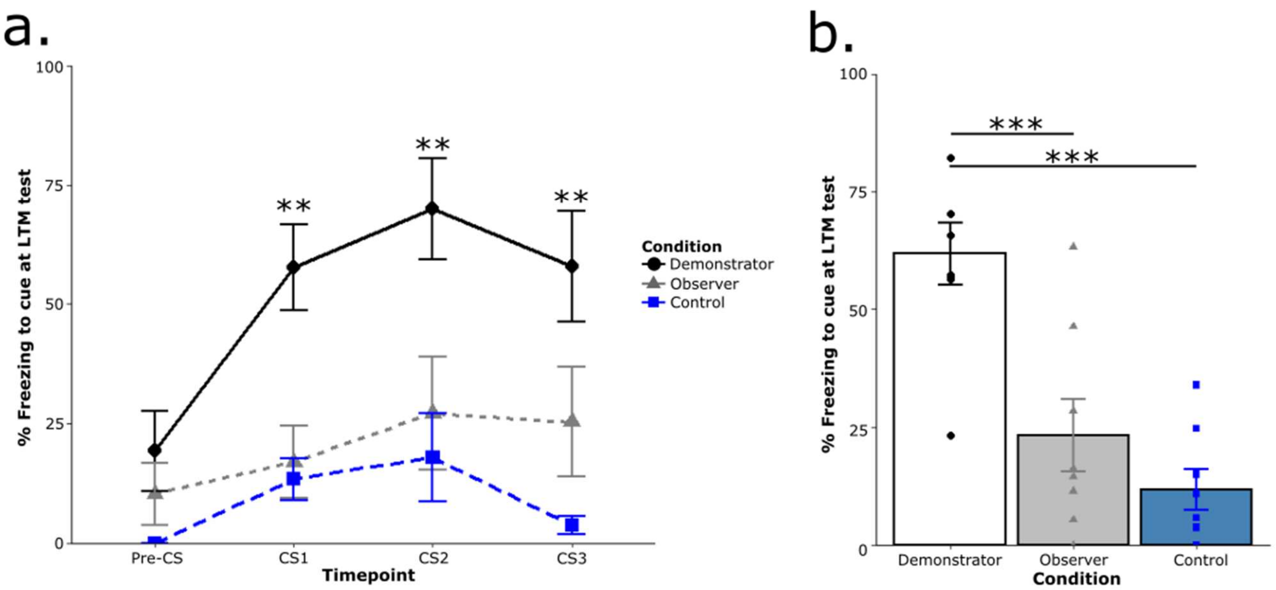


***Figure S2.*** *Dark cycle FCbP results.* (a) While demonstrators showed significantly higher freezing to the CS at all presentations as compared to observers and controls, observers did not freeze significantly more than controls and (b) this remained true when percent freezing was averaged across the three CSs.

***p < 0.001, ***p < 0.0001*

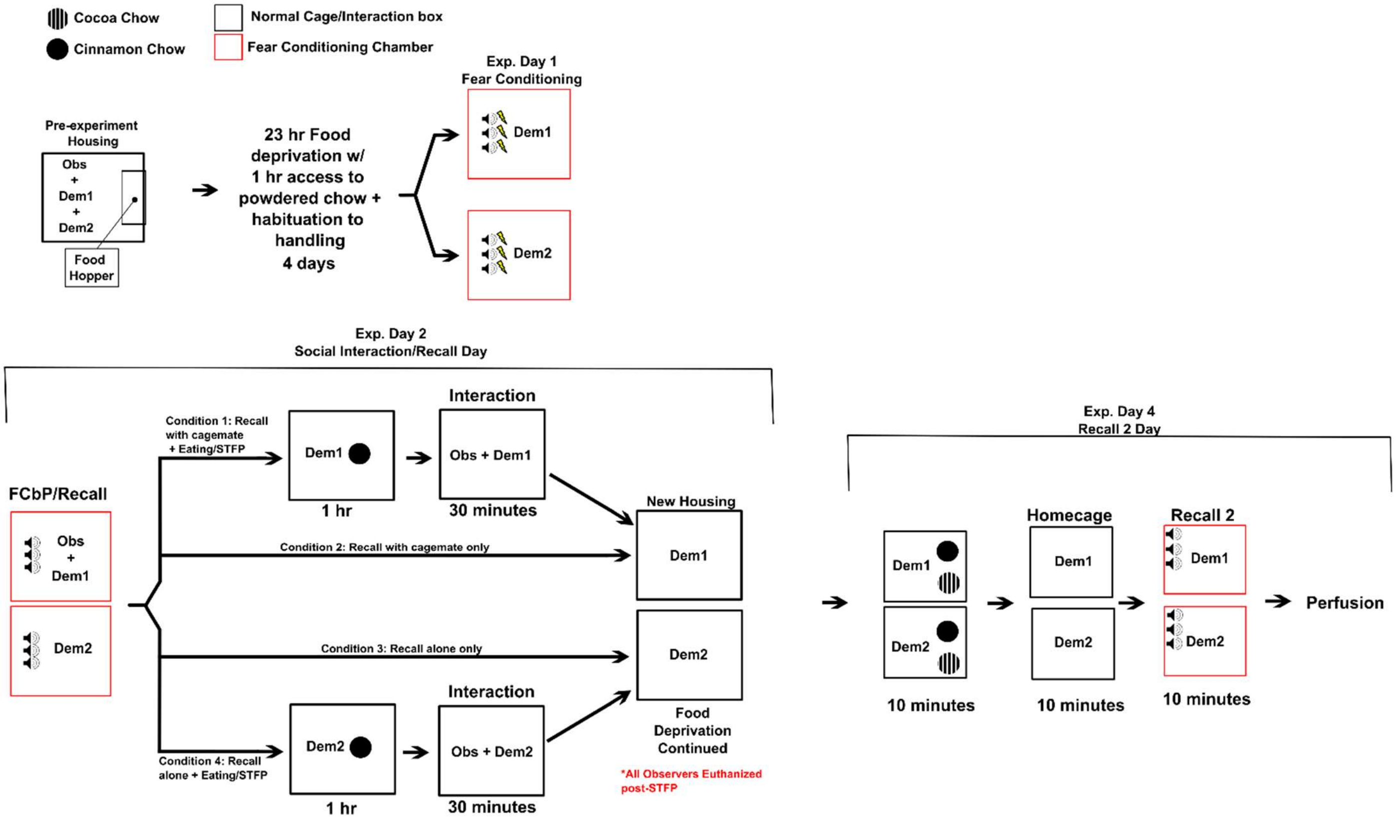


***Figure S3.*** *Effect of FCbP and STFP behavioral protocols on directly conditioned demonstrators.* This figure displays the experimental design of our follow-up experiment that was run to try and replicate the reduced freezing observed in our demonstrator animals in the primary experiment.

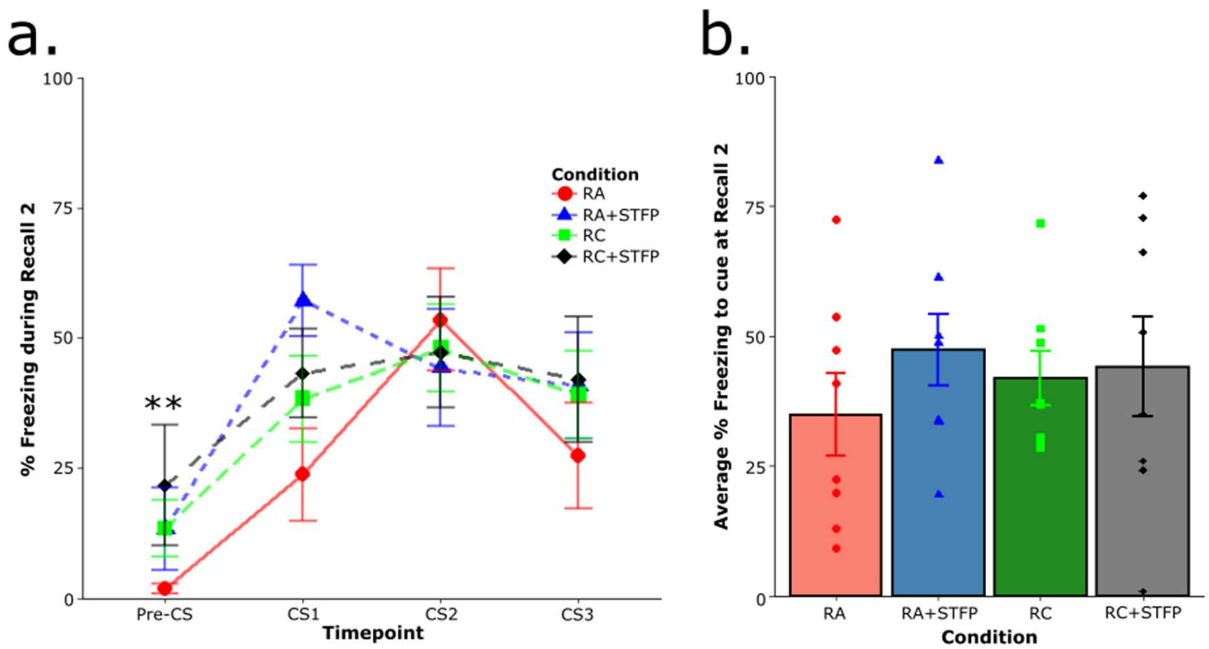


***Figure S4.*** *FCbP+STFP effect on directly conditioned rats - Results.* (a) There was no difference in freezing behavior between any of the experimental conditions (Recall alone [RA], Recall alone with STFP [RA+STFP], Recall with cagemate [RC], and Recall with cagemate and STFP [RC+STF]) and (b) this remained true when freezing was averaged across all three cues.

**Table S1. Main Effects of Gender on *Arc* Expression**

| Cell Area | Nucleus | | Cytoplasm | | Dual | |
|---|---|---|---|---|---|---|
| **Brain Region** | Statistical value | p-value | Statistical value | p-value | Statistical value | p-value |
| **lOFC** | $F_{(1,57)} = 0.31$ | 0.581 | $F_{(1,57)} = 0.01$ | 0.928 | $F_{(1,57)} = 6.18$ | 0.016* |
| **vOFC** | $F_{(1,64)} = 4.85$ | 0.031* | $F_{(1,64)} = 0.01$ | 0.905 | $F_{(1,64)} = 1.58$ | 0.214 |
| **ACC** | $F_{(1,66)} = 0.07$ | 0.788 | $F_{(1,66)} = 3.22$ | 0.078+ | $F_{(1,66)} = 15.93$ | < 0.001*** |
| **lfl** | $F_{(1,73)} = 18.05$ | < 0.001*** | $H_1 < 0.001$ | 0.98 | $F_{(1,73)} = 13.66$ | < 0.001*** |
| **vCA3** | $F_{(1,69)} = 35.47$ | < 0.001*** | $F_{(1,69)} = 60.72$ | <0.001*** | $F_{(1,69)} = 9.840$ | 0.003 ** |
| **Prl** | $F_{(1,70)} = 0.001$ | 0.982 | $H_1 = 4.3$ | 0.038* | $F_{(1,70)} = 18.11$ | < 0.001** |

*+p < 0.1, *p < 0.05, **p < 0.01, ***p < 0.001*

**Table S2. Main Effects of Condition on *Arc* Expression**

| **Cell Area** | Nucleus | | Cytoplasm | | Dual | |
|---|---|---|---|---|---|---|
| **Brain Region** | Statistical value | p-value | Statistical value | p-value | Statistical value | p-value |
| **lOFC** | $F_{(2,57)} = 0.57$ | 0.57 | $F_{(2,57)} = 0.64$ | 0.534 | $F_{(2,57)} = 0.07$ | 0.932 |
| **vOFC** | $F_{(2,64)} = 0.30$ | 0.741 | $F_{(2,64)} = 0.39$ | 0.679 | $F_{(2,64)} = 0.23$ | 0.795 |
| **ACC** | $F_{(2,66)} = 1.02$ | 0.368 | $F_{(2,66)} = 0.04$ | 0.963 | $F_{(2,66)} = 0.34$ | 0.715 |
| **IfI** | $F_{(2,73)} = 0.21$ | 0.815 | $H_2 = 0.23$ | 0.89 | $F_{(2,73)} = 1.42$ | 0.248 |
| **vCA3** | $F_{(2,69)} = 1.35$ | 0.265 | $F_{(2,69)} = 0.01$ | 0.989 | $F_{(2,69)} = 0.27$ | 0.764 |
| **PrI** | $F_{(2,70)} = 0.005$ | 0.995 | $H_2 = 2.19$ | 0.335 | $F_{(2,70)} = 0.18$ | 0.837 |

**Table S3. Condition x Gender Interactions effects on *Arc* Expression**

| **Cell Area** | Nucleus | | Cytoplasm | | Dual | |
|---|---|---|---|---|---|---|
| **Brain Region** | Statistical value | p-value | Statistical value | p-value | Statistical value | p-value |
| **lOFC** | $F_{(2,57)} = 1.595$ | 0.212 | $F_{(2,57)} = 0.125$ | 0.88 | $F_{(2,57)} = 1.08$ | 0.345 |
| **vOFC** | $F_{(2,64)} = 2.452$ | 0.094+ | $F_{(2,64)} = 1.022$ | 0.366 | $F_{(2,57)} = 1.124$ | 0.331 |
| **ACC** | $F_{(2,66)} = 0.118$ | 0.889 | $F_{(2,66)} = 0.041$ | 0.96 | $F_{(2,66)} = 0.062$ | 0.94 |
| **lfl** | $F_{(2,73)} = 0.365$ | 0.696 | $H_5 = 3.25$ | 0.662 | $F_{(2,73)} = 0.763$ | 0.47 |
| **vCA3** | $F_{(2,69)} = 0.996$ | 0.375 | $F_{(2,69)} = 0.66$ | 0.519 | $F_{(2,69)} = 0.011$ | 0.99 |
| **Prl** | $F_{(2,70)} = 3.963$ | 0.023* | $H_2 = 8.45$ | 0.133 | $F_{(2,70)} = 0.32$ | 0.727 |

*+p < 0.1, *p < 0.05*

**Table S4. Combined Food Task and Condition Effects on *Arc* Expression**

| Cell Area | Nucleus | | Cytoplasm | | Dual | |
|---|---|---|---|---|---|---|
| **Brain Region** | Statistical value | p-value | Statistical value | p-value | Statistical value | p-value |
| **lOFC** | $F_{(4,58)} = 0.567$ | 0.688 | $F_{(4,58)} = 0.818$ | 0.519 | $F_{(4,58)} = 0.233$ | 0.918 |
| **vOFC** | $F_{(4,65)} = 0.115$ | 0.98 | $F_{(4,65)} = 0.483$ | 0.748 | $F_{(4,65)} = 1.08$ | 0.376 |
| **ACC** | $F_{(4,67)} = 0.796$ | 0.53 | $F_{(4,67)} = 0.12$ | 0.975 | $F_{(4,67)} = 0.426$ | 0.79 |
| **lfl** | $F_{(4,74)} = 0.116$ | 0.977 | $F_{(4,74)} = 0.24$ | 0.915 | $H_{(4)} = 4.85$ | 0.304 |
| **vCA3** | $F_{(4,70)} = 1.51$ | 0.209 | $F_{(4,70)} = 0.653$ | 0.627 | $F_{(4,70)} = 0.729$ | 0.575 |
| **PrI** | $F_{(4,70)} = 0.377$ | 0.824 | $H_4 = 2.27$ | 0.686 | $F_{(4,70)} = 1.120$ | 0.354 |
